# Shared and host-specific transcriptomic response of *Pseudomonas donghuensis* P482 to the exudates of tomato (Dicot) and maize (Monocot) shed light on the host-adaptive traits in the promiscuous root colonizing bacteria

**DOI:** 10.1101/2022.07.13.499613

**Authors:** Dorota M. Krzyżanowska, Magdalena Jabłońska, Zbigniew Kaczyński, Małgorzata Czerwicka-Pach, Katarzyna Macur, Sylwia Jafra

## Abstract

Plants of different genotypes and physiological states recruit different populations of root microbiota. The selection is driven by the immune response of the plant and the composition of root exudates. Some bacteria, including *Pseudomonas* spp., are promiscuous root colonizers. It is yet unclear what particular changes in lifestyle enable them to thrive in the company of different plant hosts. In this study, we used RNAseq to identify genes of the differential (host-specific) and shared (host-independent) transcriptomic responses of a biocontrol strain *Pseudomonas donghuensis* P482 to the root exudates of two phylogenetically distinct plant species, tomato (Dicot) and maize (Monocot), both of which can be colonized by the bacterium. The host-independent response of P482 to exudates involved upregulated expression of arsenic resistance genes and bacterioferritin synthesis. Contrary, we observed downregulation of pathways related to sulfur assimilation, sensing of ferric citrate and/or other iron carriers, the acquisition of heme, the assembly of the type VI secretion system, and amino acid transport. Pathways upregulated in P482 specifically by tomato exudates included nitric oxide detoxification, repair of iron-sulfur clusters, respiration through the cyanide-insensitive cytochrome *bd*, and catabolism of amino acids and/or fatty acids. The maize-specific response included upregulation of genes associated with motility, the activity of MexE and two other RND efflux pumps, and copper tolerance. To provide more context to the study, we determined the chemical composition of exudates by GC-MS, NMR, and LC-SRM. Our results bring new insight into the host-driven metabolic adaptations of promiscuous root colonizing bacteria.

**Significance statement:** Understanding factors determining the composition and the activity of plant-associated microbiota is crucial to harnessing their potential to benefit plant health. Traits that enable microorganisms to colonize plants have long been the subject of study, with many important factors identified for particular host-microbe systems. However, studies involving more than a single plant host are rare. This results in many unanswered questions on the host-specific and universal aspects of metabolism that enable more promiscuous root colonizers to interact with different plant hosts. The presented study begins to fill this knowledge gap by providing data on the metabolic pathways involved in the differential and shared response of *Pseudomonas donghuensis* strain P482 to the exudates of phylogenetically distant plant species: tomato and maize.

## Introduction

Every plant in nature is accompanied by a diverse community of microorganisms. Rhizosphere is a hotspot of microbial activity due to the presence of nutrients secreted by the plant through the roots (1). Root exudates comprise of primary metabolites, such as organic acids, amino acids, and sugars, as well as secondary metabolites with bioactive or signaling properties (reviewed in (2)). The composition of root exudates varies from one plant species to another. Moreover, it is affected by the developmental stage of the plant, nutrient availability, and other factors affecting the plant’s physiological status (3). Those differences, together with the plant’s innate immunity, act as a filter shaping the composition and the activity of the root microbiota (4) (5, 6).

From the microbial end of the plant-microbe interaction, securing a place in the rhizosphere requires the ability to colonize the roots of the plant host, evade plant defenses, use carbon sources that the host provides (with the capacity for quick adjustments), and efficiently compete with other microorganisms. These requirements are perfectly met by the bacteria of the genus *Pseudomonas.* Pseudomonads can be found in versatile ecological niches, however many live in association with plants. The ubiquity of these microorganisms can be attributed to efficient adaptation of the cells to the fluctuations in the environment and their potential to metabolize multiple carbon sources (7, 8). The competitive edge of pseudomonads is further enhanced by production of a wide range of secondary metabolites, including antimicrobials and iron-scavenging compounds (9). Although genus *Pseudomonas* includes animal and plant pathogens, with the respective examples of *P. aeruginosa* and *P. syringe*, most pseudomonads are either harmless commensals or beneficial organisms (10). Many strains are studied for plant growth promotion, alleviation of abiotic stress in plants or biocontrol against pathogens (11, 12). Among them there is *Pseudomonas donghuensis* P482, an isolate from the rhizosphere of tomato (*Solanum lycopersicum* L.) (13). Due to the production of secondary metabolites and volatile organic compounds, the strain is able to inhibit the growth of several bacterial and fungal plant pathogens, including soft rot *Pectobacteriaceae* (SRP) (former *Erwinia* spp.), *Pseudomonas syringae*, *Rhizoctonia solani*, *Fusarium culmorum, Verticillium dahliae* and the oomycete *Phytium ultimum* (14) (13, 15) .

There are no comprehensive studies regarding the spectrum of plant hosts that certain *Pseudomonas* strains can colonize, and with what efficiency. However, reports of useing certain pseudomonads as biocontrol agents on a different plant than the plant of origin, or on more than a single crop, suggest that pseudomonads are rather promiscuous regarding the plant hosts, with some strains reported to be able to switch their lifestyle from the plant-associated to associated with humans or animals (16, 17). Strain *P. donghuensis* P482, originating from the rhizosphere of tomato, was shown to colonize the rhizosphere of soil-grown potato (18). We also observed its potential to colonize the roots of Monocot plants in small scale *in vitro* experiments (unpublished), encouraging us to perform soil-based colonization experiments in this study .

The concept behind the assembly of plant microbiome is streamlining, that is possibility of a particular root zone to be inhabited by a limited variety of species (19). It was also shown that more phylogenetically distant plant hosts recruit more distinct bacterial communities (20). Therefore, host promiscuity of some microorganisms, including pseudomonads, raises questions on both the specific and the universal aspects of microbial metabolism involved in the interaction with different plants. This question, important from the perspective of microbiome assembly and functioning, is difficult to answer based on a meta-analysis of existing data. Scientific methods relying on next generation sequencing, including RNAseq, greatly improved our ablitity to collect vast datasets. However, most studies involve single plant-microorganism setups.

In this work, we provide new insight into the specific and universal aspects of interaction between pseudomonads and different plant hots by identifying genes of the differentiating (‘host-specific’) and shared (‘universal’) transcriptomic response of strain *Pseudomonas donghuensis* P482 to the root exudates of two phylogenetically distinct plant species: tomato (Dicot) and maize (Monocot).

## Materials and Methods

### Bacterial strains and culture conditions

Bacterial strains used in this study included *Pseudomonas donghuensis* P482, an isolate from the rhizosphere of tomato (13, 21), and its spontaneous rifampicin resistant mutant P482 Rif (18). Routine propagation of bacteria was performed at 28 °C in Miller’s LB liquid medium (Roth), with agitation (120 rpm), or on solid media plates with LB or TSB (Oxoid) supplemented with agar (15 g L^-1^). Experiments requiring cultivation of P482 in the presence of root exudates were performed in a single carbon source medium (1C medium) comprising: basic M9 salts (MP Biomedicals), 10 mM of Mg_2_SO_4_, 0.5 mM CaCl_2_ (22) and 0.4% glucose. Where indicated, rifampicin (50 µg mL^-1^) (Sigma-Aldrich/Merck), ampicillin (50 µg mL^-1^) (Polfa Tarchonin) and cyclocheximide (100 µg mL^-1^) (Sigma-Aldrich/Merck) were supplemented to the media as selective agents.

### Plant material

Seeds of tomato (*Solanum lycopersicum* L.) cv. Saint Pierre (Vilmorin Garden, Poland) were purchased in a gardening store. Seeds of maize (*Zea mays* L.) cv. Bejm were purchased from the Plant Breeding and Acclimatization Institute (Ihar; Smolice, Poland). The same batch of seeds was used in all experiments per each species.

### Colonization of roots of soil-grown plants

Plant seeds were germinated for 4 days in glass Petri dishes lined with Whatman filter paper. The filter paper was moistened with distilled water and the plates were sealed with parafilm to retain humidity. Tomato was germinated in the dark at 24 °C and maize was germinated in light at 22 °C.

To obtain bacterial inoculum, strain P482 Rif was cultured for 24 h on TSB agar supplemented with rifampicin (50 µg mL^-1^). Cells were harvested, suspended in sterile saline and the turbidity of the suspension was adjusted to 6 units in McFarland scale (10^9^ colony forming units per mL; CFU mL^-1^). The cells were harvested by centrifugation (4 min, 3800 RCF). The pellet was re-suspended in 1% carboxymethylcellulose (Calbiochem, Germany), in a volume equal to the one of the discarded supernatant to maintain the established cell density. The germinated seeds were coated in the inoculant and potted into the soil to 12 cm dia. pots. Tomato was grown in universal potting soil (Kronen) and maize was grown in a mixture of potting soil and silica sand (1:1, v/v) to assure proper drainage of the substrate. Seven pots were prepared per plant species, with one seed per pot for maize and three seeds per pot in the case of tomato. Plants were grown for 14 days in a phytotron chamber, 16/8 h day/night photoperiod, 22 °C, 60% humidity, white light (Philips Master TL-D 36W/840 5B, Philips).

Following 14 days of growth in soil (18 days in total, including germination), plants were harvested. Roots were shaken to remove the unbound soil, and the rhizosphere samples were placed in universal filter extraction bags (Bioreba). Near complete rhizosphere systems were collected to avoid error resulting from variation in the special distribution of bacterial cells on the root. Nine mL of sterile saline was added to an extraction bag per 1 g of sample. The bag was sealed and plant material was homogenized using a hammer. Immediately post homogenization, each suspension was serially diluted (1:9). Ten µL aliquots of each dilution were spot plated on a selective medium, three technical replicates each, as described previously for high-throughput plating of bacterial formulations (23). The applied medium was TSB agar supplemented with rifampicin (50 µg mL^-1^) and ampicillin (50 µg mL^-1^), added to select for P482 Rif, and cycloheximide (100 µg mL^-1^) to inhibit the growth of soil fungi. Plates were incubated at 28 °C for up to 48 hours. Colony count enabled calculation of the titer of P482 Rif per gram of fresh weight of the rhizosphere samples. The experiment was performed twice (experiments 1 and 2; E1 and E2).

Data from both experiments (E1 and E2) were pooled for statistical analysis to obtain 14 data points per species. The normality of data was determined using Shapiro-Wilk’s test. Non-parametric two-tailed Mann–Whitney U test (α=0.05) was applied to determine the significance of differences between groups, with the help of an online tool available at http://www.statskingdom.com/.

### Plant growth in gnotobiotic conditions Surface sterilization of seeds

Seeds of tomato were sterilized in 2 mL tubes, ca. 50 seeds per tube. Sterilization protocol was as follows: 1 min in 1 mL of 70% ethanol – gentle mixing by tube inversion, 3 min vortexing in 1 mL 3% NaOCl (Chloraxid, Cerkamed, Polska), rinsing 3 times with 1 mL of sterile distilled water – mixing by tube inversion. Seeds of maize were surface sterilized in 50 mL conical tubes, ca. 20 seeds per tube. The protocol was as follows: 2 times incubation, 15 min. each, in 35 mL of 3% NaOCl, followed by 10 min in 35 mL 70% etanol and 3 times 5 min rinsing in sterile distilled water. For all incubation steps concerning the sterilization of maize seeds, the tubes were fixed flat in a rotary shaker (150 RPM).

### Seed germination

Seeds of both tomato and maize were transferred aseptically to Petri plates (dia. 10 cm) with Germination Medium (GM) (0.5 × Murashige and Skoog Medium including Gamborg B5 Vitamins (Duchefa Biochemie), 2% sucrose (Chempur), 0.2% wheat peptone (Sigma-Aldrich/Merck) and 0.7% of plant agar (Duchefa Biochemie), pH ≈6.1). The GM was developed and applied in this work as a medium that supports the growth of microorganisms and therefore enables verification of the efficiency of the sterilization process. The seeds of both plants were germinated for five days. Tomato was germinated in the dark at 24 °C, 12 seeds per plate. Maize seeds were germinated in light (16/8 h day/night photoperiod) at 22 °C in a plant growth cabinet (Sanwood), 6 seeds per plate. To exclude contamination, each plate with microbial growth detected for even a single germinated seed was discarded.

### Plant culture in sterile gravel

The germinated seeds were moved aseptically to grow in sterile quarts gravel caliber 1.4-2 mm (AQUAEL, Poland). Maize was cultured in ¼ Hoagland’s medium (Hoagland’s No. 2 Basal Salt Mixture, Sigma-Aldrich/Merck), pH adjusted to 5.6-5.8 with KOH, in 900 mL glass jars with screw caps equipped with cotton wool corks, 3 seedlings per jar. Tomato was cultured in ½ Hoagland’s medium in GA-7 Magenta vessels (Sigma-Aldrich/Merck), 6 plants per vessel. To prepare the setup, quartz gravel was washed 3 times in distilled water, transferred to aluminium baking forms and dried in an oven (≥1 h at 140 °C). After cooling, 200 g of gravel was weighted to either the GA-7 Magenta vessels or the 900 mL glass jars. The vessels were autoclaved together with the gravel. Sterile Hoagland’s medium was aliquoted aseptically, 50 mL per growth vessel. Germinated seeds were transferred to the containers, six seeds per container in the case of tomato and three seeds for maize. Containers to which no plants were potted were prepared as controls. The plants were grown in a phytotron chamber at 22 °C, 16 h light/8 h dark photoperiod, white light (Philips Master TL-D 36W/840 5B, Philips) for 13 days. Both plant species established a second pair of leaves within this period (picture in SI Appendix, Fig. S1). Each vessel was sampled for sterility six days before harvest by plating 100 µL of the aspired media culture on LA plates. Vessels for which microbial growth was observed after 48 hours at 28 °C were discarded.

### Collection of root exudates

The exudates were collected in high-purity water (Purelab Classic, Elga). All procedures were performed in sterile conditions. Plants grown in gnotobiotic conditions were gently removed from the gravel. To avoid root damage, the gravel was flooded with ca. 50 mL of water to loosen up the particles. Plant roots were placed in water for 2-4 minutes to wash off the remaining medium and the potentially damaged cells. Next, the plants were transferred to glass beakers containing a fresh aliquot of sterile water: 75 mL (in 250 mL beakers) per 30 plants in the case of tomato and 100 mL (in 1000 mL beakers) per 6-7 plants for maize. The beakers were secured with aluminium foil and incubated in a phytotron chamber for precisely two hours. For the small tomato plants, some leaves were touching the water surface. In total, exudates from 174 tomato plants and 48 maize plants were collected. The samples from each species were pooled and filter-sterilized through a 0.22 µm low-binding PES membrane using Rapid Flow Rapid-Flow Sterile Disposable Filter Units (Nalgene/Thermo Scientific). The samples were frozen at −80 °C and freeze dried. Chemical analyses required sample pretreatment prior to lyophilization (details provided in the respective sections).

### Chemical analysis of the composition of root exudates

The chemical composition of the root exudates was assessed by gas chromatography mass spectrometry (GC-MS), nuclear magnetic resonance (NMR) and, specifically for the relative quantity of amino acids, with liquid chromatoraphy-selected reaction monitoring mass spectrometry (LC-SRM). Details of the experimental procedures are provided in Supplementary Information (SI Appendix).

### Growth of P482 in the presence of root exudates

The growth rate of P482 in root exudates was evaluated for cultures in the 1C medium supplemented with three different concentrations of the dry weight of either tomato or maize root exudates: 0.02 mg L^-1^, 0.1 mg L^-1^, 0.2 mg L^-1^. In addition, growth in the 1C medium alone was measured for reference. The experiment was performed in a 96-well plate format. Two hundred µL of 1C medium, with or without the root exudates, was inoculated with 4 µL (50:1) of the starter culture, four technical replicates per combination. The starter culture for the experiment was an overnight culture of P482 in 1C medium with turbidity adjusted with fresh medium to 0.8 units in McFarland scale. The inoculated 96-well plate was sealed with foil (Sarstedt) and grown in EPOCH microplate reader (BioTek) at 28 °C, temperature gradient between the lid and bottom set to 1 °C to prevent condensation, orbital shaking, and with OD_600_ measurements taken each 20 minutes for a total of 43 hours. The experiment was conducted twice.

Additionally, the pH of the 1C medium, with and without supplementation with root exudates (0.2 mg L^-1^), was measured for sterile solutions (initial pH) and following growth of P482 to an early stationary phase (post-culture pH). The measurement was performed with MQuant pH indicator strips, pH range 2-9, resolution 0.5 unit (Supelco).

### Harvest of bacterial cells for gene expression analyses

To obtain bacterial cells for the isolation of RNA, the P482 was cultured in 96 well plates, according to the protocol described above for the growth in root exudates. The strain was cultured in 1C medium alone (reference) or 1C medium supplemented with either maize or tomato root exudates (0.2 mg L^-1^ exudate dry weight). To avoid collecting cells at different growth phases, the growth rate in tested media was monitored in real-time. Bacterial cells were harvested upon reaching the early stationary phase, following 33 h of growth in unsupplemented 1C medium and 24 and 28 hours, respectively, for the growth in 1C medium with maize or tomato exudates. The cells were harvested from 5-6 wells and pooled to obtain a single sample containing ca. 10^9^ of cells. In total, 3 pools were obtained per treatment, adding up to a total of 9 samples processed for the isolation of RNA. In order to harvest the cells, the plate was removed from the microplate reader for up to 5 minutes, the foil was cleansed with 70% ethanol and a cut was made over each target well with a sterile scalpel. Cultures were aspired from the wells. P482 cells were immediately harvested by centrifugation (2 min, 10000 RCF), re-suspended in 250 µL of fixRNA (Eurx) and pelleted again. The fixed pellets were kept frozen at −80 °C until further processing.

### Isolation of RNA and cDNA synthesis

Fixed bacterial pellets (ca. 10^9^ of cells per sample) were thawed on ice. To remove the excess of fixing agent, 200 µL of sterile TE buffer (Eurx) was added atop the pellet (no pipetting), the samples were centrifuged (3 min., 5000 RCF) and the supernatant was aspired. The RNA was isolated from the cell pellets using RNeasy Mini Kit (Qiagen) according to the manufacturer’s protocol involving lysozyme and proteinase K treatments. Optional on-column DNase digestion was performed using DNaseI from an alternative supplier (Eurx). Samples were eluted with DEPC-treated water (Eurx). To eliminate potential carryover of gDNA, samples were treated with TURBO DNA-free™ Kit (Thermo Fisher Scientific). Purified RNA was stored at −80 °C in aliquots (to avoid freeze-thaw cycles). The concentration of RNA was measured using NanoDrop 2000 (Thermo Fisher Scientific) and the integrity of samples was evaluated using agarose gel electrophoresis. The lack of gDNA contamination in the RNA samples was confirmed by qPCR with primers targeting *gyrB* of strain P482 (for details, see the cDNA synthesis and qPCR section below).

### RNAseq and data analysis

Procedures involving quality control of RNA samples, depletion of rRNA, library preparation, sequencing with Illumina technology and transcriptome assembly was performed at Baseclear (Leiden, The Netherlands). Three independent samples were processed for P482 grown under each of the three tested conditions: 1C medium with tomato root exudates (0.2 mg L^-1^), 1C medium with maize exudates (0.2 mg L^-1^) and 1C medium alone (9 samples in total).

RNA isolated from all samples passed the quality check regarding concentration and integrity. Depletion of rRNA proved challenging at the time of the experiment – the Illumina’s Ribo-Zero rRNA Removal Kit, a method of choice for this procedure in many commercial sequencing facilities, was unexpectedly withdrawn from the market. As an alternative, the samples we treated with MICROBExpress Bacterial mRNA Enrichment Kit (Thermo Fisher Scientific). Although the set was shown efficient for the related *Pseudomonas putida*, in the case of *P. donghuensis* P482 it provided only partial depletion of rRNA. We successfully dealt with this obstacle and obtained sufficient coverage by increasing the minimal sequencing depth from the 3 GB to 5 GB per sample. The obtained RNA-seq reads, following demultiplexing and filtering, were mapped to the reference genome of P482 (Genbank: JHTS00000000.1) (24) using STAR aligner version 2.7.1 (25).

Further data analysis involved differential gene expression performed using DESeq2 version 1.22.2 (26). The compared groups included P482 grown: 1) in the presence of tomato root exudates vs in 1C medium alone; 2) in the presence of exudates of maize vs in 1C medium alone and 3) in the presence of root exudates of tomato vs in the presence of the exudates of maize. Principal component analysis (PCA) was applied to verify wheather the replicate samples form separate clusters. Listing all differentially expressed genes, including calculation of log_2_ changes in expression and p-values and assignment of functional categories, was part of the outsourced bioinformatic service (Baseclear, The Netherlands). Further analyses were performed in-house. The company-provided data were filtered using cutoffs of 1.5 log_2_ fold change (log_2_FC) in gene expression for biological significance and α=0.05 (p<0.05). The applied threshold of biological significance is one of the lowest applied in RNAseq analyses, and was chosen to minimize the risks of enriched pathways or networks not being identified in downstream analysis due to the excluded genes. Subsequently, overlapping groups of differentially expressed genes were determined and visualized with the help of BioVenn (27). Proteins were assigned to Clusters of Orthologous Groups (COGs) using eggNOG mapper 5.0 (http://eggnog-mapper.embl.de/) (28) and to KEGG metabolic pathways using BlastKOALA (29). Enrichment within COGs and KEGG pathways was established using the genome of P482 as reference (Genbank WGS accession: JHTS00000000.1),with Fisher’s exact test applied to determine statistical significance of differences in the proportion of categories. Networking between genes and cluster enrichment was analyzed using STRING version 11.5 (September 2021) (https://string-db.org/) (30), with all types of interaction sources enabled (text mining, experiments, databases, co-expression, neighbourhood, gene fusion, co-occurrence). Additionally, we investigated enrichment in functional domains as possible by this tool (based on pfam, SMART and InterPro databases, and ‘STRING clusters’). The genome *P. donghuensis* HYS (31) was used as a reference genome for the STRING analysis. HYS, established to be a sibling strain of P482 (21), was the most closely related organism available in the STRING database. Additionally, ranking lists were created to pinpoint the most upregulated and downregulated genes in the sets of interest.

### cDNA synthesis and real-time qPCR

A fraction of each RNA sample analyzed by RNAseq was reverse-transcribed to cDNA using Transcriptor First Strand cDNA Synthesis Kit (Roche) and random hexamer primers. The amount of RNA per reaction was adjusted to 500 ng. The optional matrix denaturation step was applied according to the manufacturer’s protocol. cDNA was stored at −20 °C prior to further processing.

Real-time qPCR was carried out in a 96-well plate format using the CFX96 instrument (Bio-Rad), using a protocol modified from (32). Each reaction mixture (V_t_=20 μL) comprised of 2× Power SYBR Green Mastermix (Thermo Scientific, USA), forward and reverse primers (300 nM each) and 4 μL of 1:7 diluted cDNA. The cycling conditions were as follows: 95 °C for 10 min, 40 cycles of 95 °C for 15 s and 60 °C for 1 min, with a final melt curve 55-95 °C (0.5 °C/ 5 s increment). Sample maximization design was applied so that all samples for each target were tested in a single run. Primers were designed using Primer3Plus (33). For primer details, refer to SI Appendix, Table S1. Primer efficiencies was calculated based on 7-point standard curves for which 10-fold serial dilutions of post-PCR products were used as templates. The size of the amplicons was confirmed by gel electrophoresis (SI Appendix, Fig. S2). Gene expression analysis was performed using qbase + (34). The *gyrB* and *rpoD*, two reference genes used for data normalization, were chosen based on (32) and confirmed for stability in the analyzed dataset (average M=0.543; CV=0.181). Expression was scaled to the samples originating from P482 grown in 1C medium without exudates (untreated). Results expressed as Calibrated Normalized Relative Quantity (CNRQ) were log_2_-transformad to obtain fold change. Statistical significance of differences in expression was evaluated using the unpaired t-test.

**Table 1.**
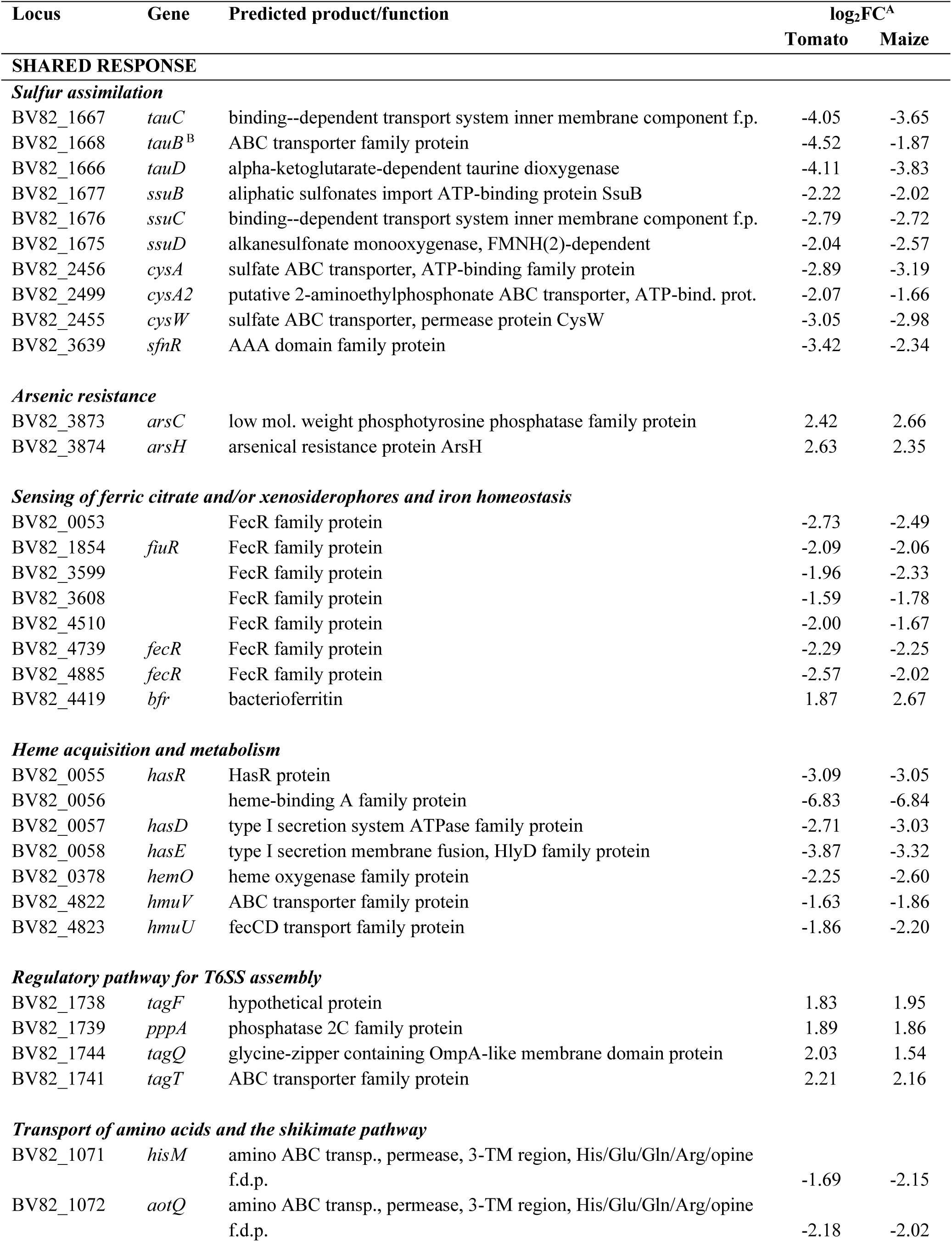
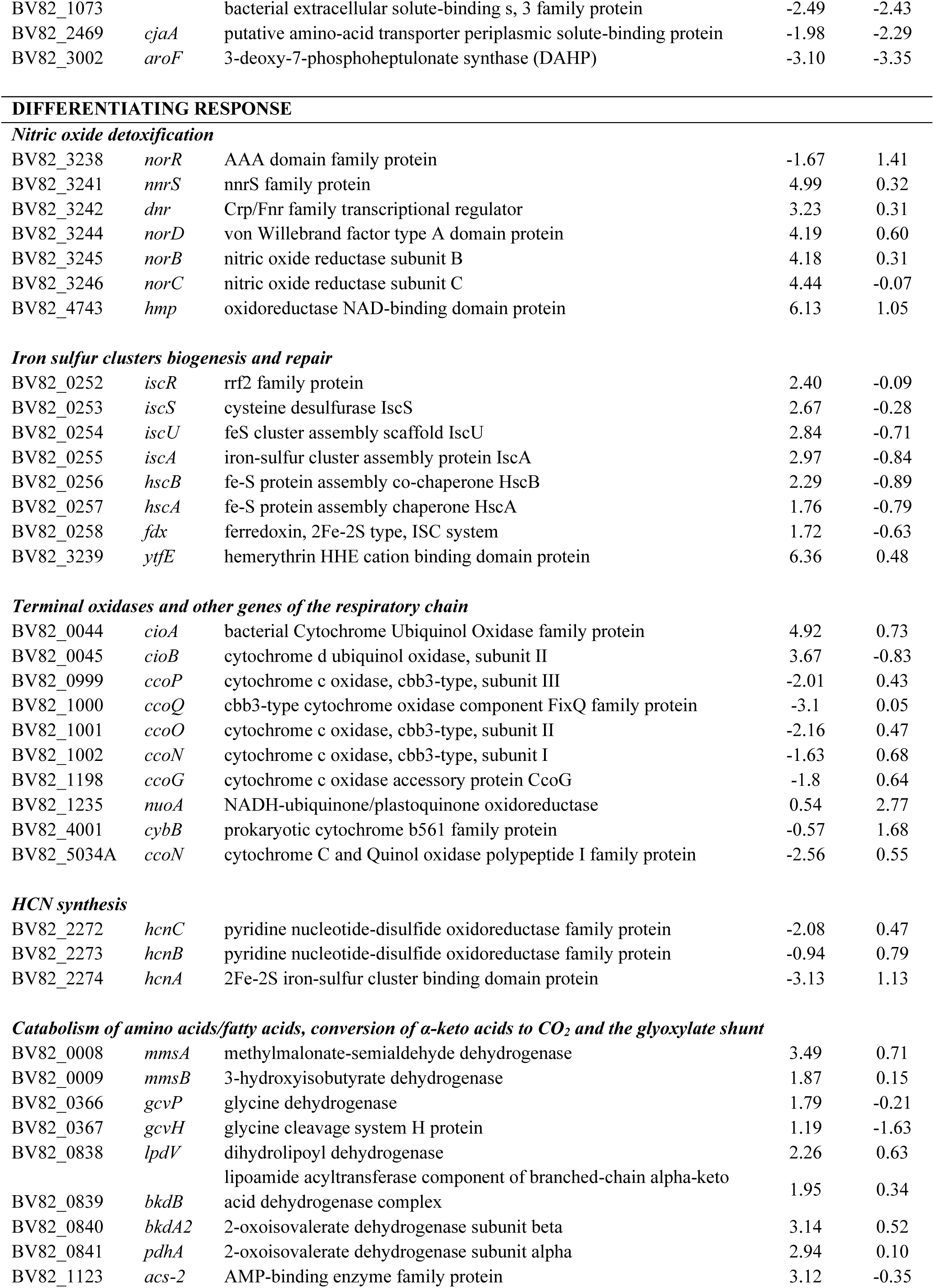
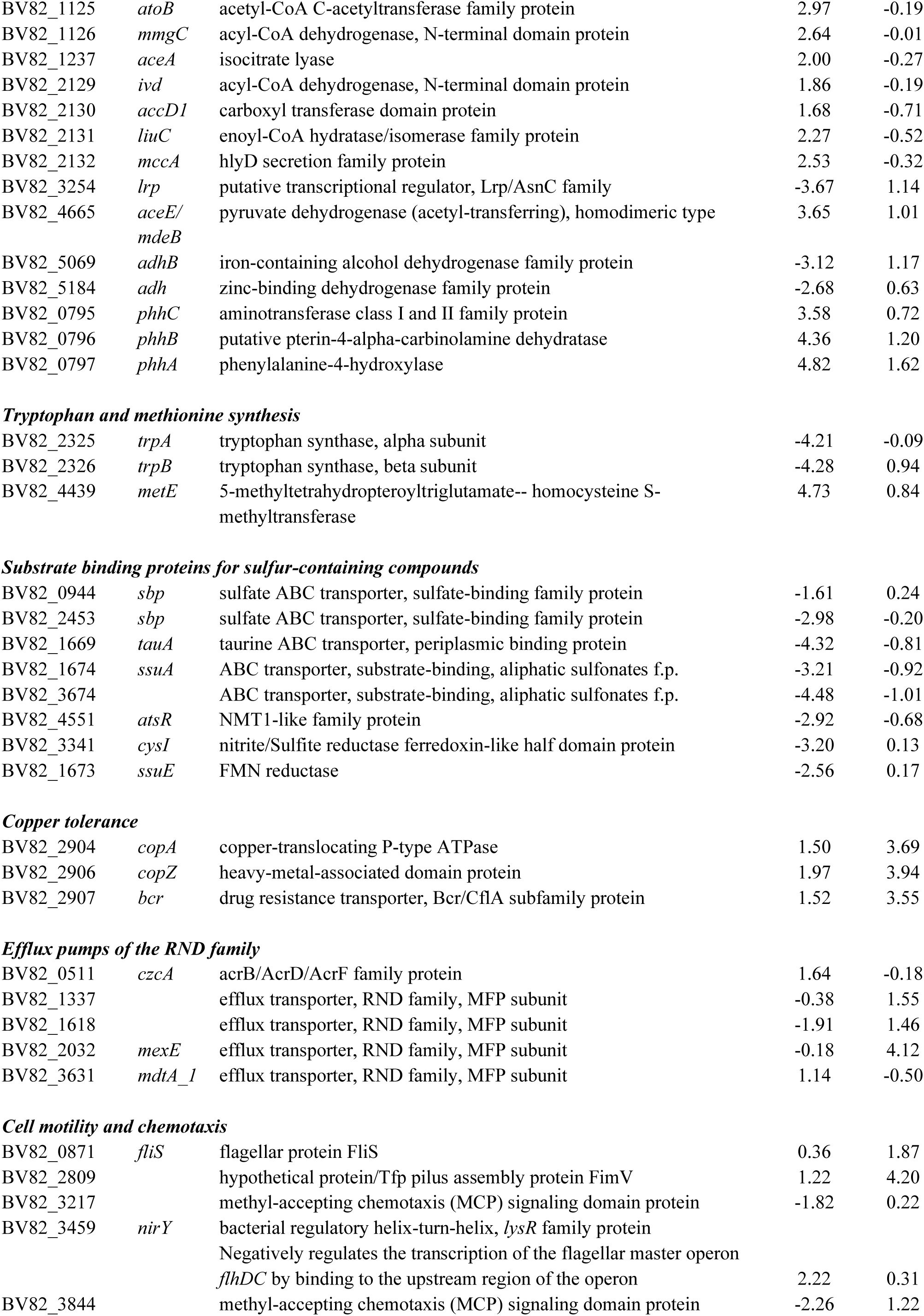
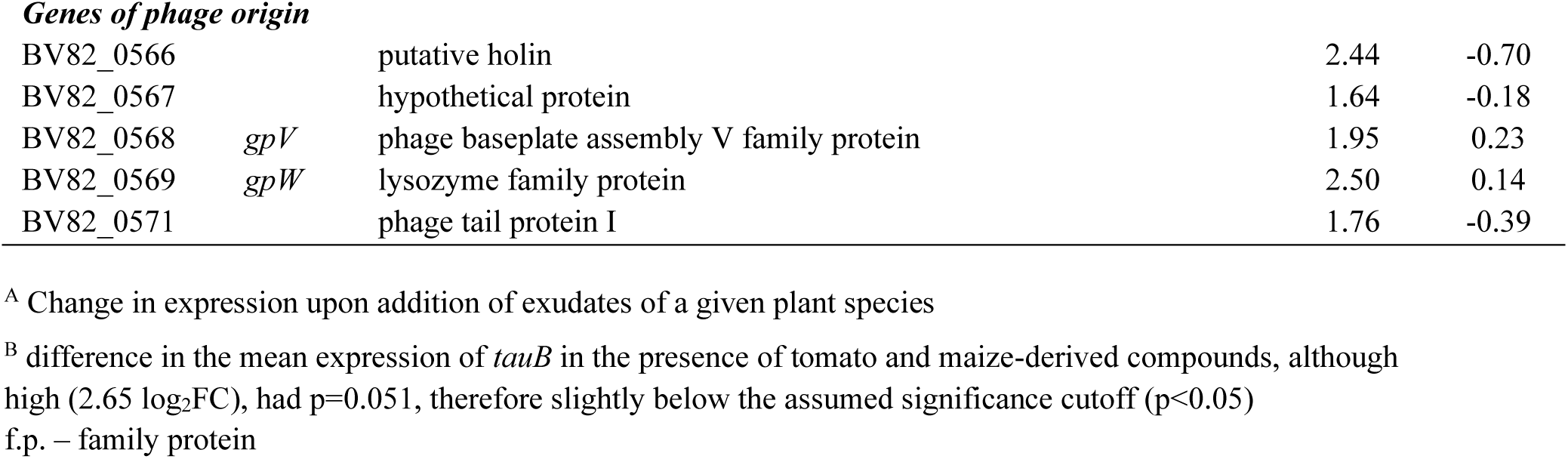
Selected genes of the shared and differentiating responses of P482 to plant exudates.

## Results

### Colonization of soil-grown tomato and maize by strain P482

The ability of *P. donghuensis* P482 to colonize the roots of tomato and maize was assessed using P482 Rif. The size of the bacterial population was established on the roots of 18-day old plants. As shown by the titer of P482 Rif per gram of rhizosphere sample, the strain efficiently colonises the rhizosphere of both plant species (Fig. 1). Under the applied experimental conditions, the titer of P482 Rif on the roots of tomato was approx. 1.5 orders of magnitude higher than that on the roots of maize, with a median value of 8.1 × 10^6^ CFU g^-1^ (and the average 9.8 × 10^6^ CFU g^-1^) in the case of tomato and a median of 3.1 × 10^5^ CFU g^-1^ (and the average 1.3 × 10^6^ CFU g^-1^) in the case of maize. The difference was statistically significant (p=0.000007). On the contrary, no significant difference was found between independent experiments (1 and 2), for both plant species, confirming the consistency of data. CFU counts obtained for all processed samples (raw data) are available in SI Appendix, Table S2.

**Fig. 1.**
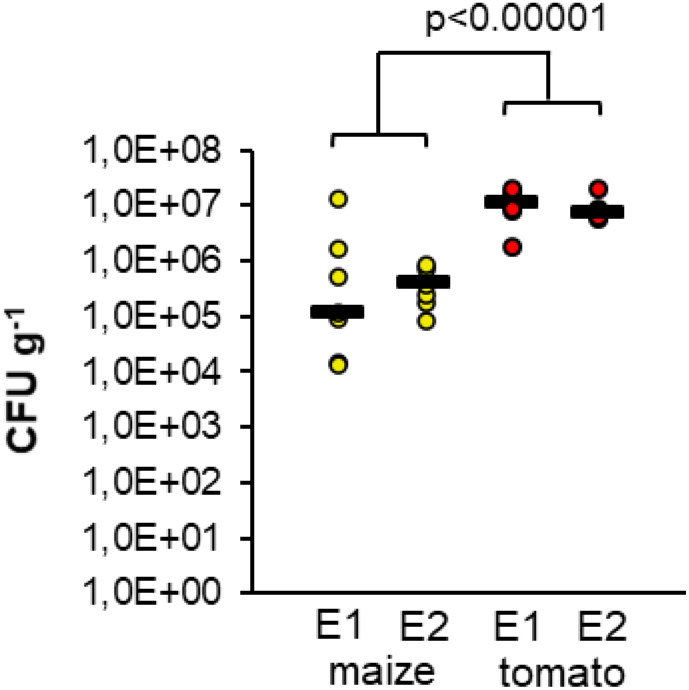
The population size of *P. donghuensis* P482 Rif in the rhizosphere of soil-grown tomato and maize. Each data point represent the titer of cells per gram of rhizosphere sample (CFU g^-1^). Horizontal lines on the chart show median values. E1 and E2 stand for independent experiments, each including 7 replicates per species. Statistical significance of differences observed between tomato and maize, expressed by p-value, was calculated for data pooled from E1 and E2 (14 datapoints in total) using the Mann–Whitney U test (α=0.05).

### Growth of P482 in 1C medium supplemented with different concentrations of root exudates

The influence of root exudates on the growth rate of P482 was evaluated for three different concentrations of exudate dry weight: 0.02 mg L^-1^, 0.1 mg L^-1^ and 0.2 mg L^-1^. When added to the 1C medium, neither exudates of tomato nor maize showed a negative effect on the growth rate of the strain. Moreover, the addition of exudates stimulated the growth of P482 in a concentration-dependent manner (SI Appendix, Fig. S3 A, B). When compared to an unsupplemented medium, the cultures grown in the presence of exudates reached the stationary phase faster and achieved higher end-point cell density. The effect was observed for all three tested concentrations of maize exudates and in all but the lowest concentration in the case of the tomato exudates (0.02 mg L^-1^), where the growth characteristics were similar as in the 1C medium alone (SI Appendix, Fig. S3 A, B).

For the RNAseq, we supplemented the cultures of P482 with the highest of the tested concentrations of exudates. This strategy was designed to increase the chances of seeing changes in exudate-induced gene expression. Bacterial cells from all treatments were harvested at the early stationary phase, that is, following 33 h of growth in unsupplemented 1C medium and 24 and 28 hours for the growth in the same medium with maize or tomato exudates, respectively (SI Appendix, Fig. S3 C).

pH measurements were taken for unsuplemented 1C medium and 1C supplementated with either tomato or maize exudates (0.2 mg L^-1^). No difference was observed between the three combinations. The initial pH of all samples equaled 7, with a decrease to pH 6 following the growth of P482 to early stationary phase.

### RNA sequencing and mapping of transcripts

Sequencing yield for the nine processed samples varied from 5.84 to 9.28 Gbp, translating to 20.8 to 31.3 million of paired-end reads, respectively (SI Appendix, Table S3). The fraction of multimapped reads ranged from 63.31-72.93% due to incomplete digestion of

rRNA. This was compensated by the applied sequencing depth, yielding above 2 million non-rRNA fragments mapping to single locations in all but a single sample, with an average of 4 million (SI Appendix, Table S4). In total, the performed experiment enabled the detection of expression of 5168 genes, comprising 99% of all ORFs in the version of the P482 genome used as reference. The principal component analysis confirmed that data points representing transcription profiles of biological replicates cluster together, while separately for the analyzed growth conditions (SI Appendix, Fig. S4).

### Differential gene expression analysis and grouping of genes into subsets

Differential gene expression analysis was performed to find differences between transcription profiles of P482 grown in the presence of tomato or maize exudates compared to unsupplemented 1C medium (background). Genes expressed differently compared to the background were termed the differentially expressed genes (DEGs). Terms tiDEGs and miDEGs were introduced to distinguish between the tomato-induced and maize-induced DEGs, respectively.

Among 5168 genes detected in the transcriptome of P482, a significant change in expression in response to tomato-derived compounds was observed for 440 genes (8.51% of the transcriptome; 62% upregulated, 38% downregulated). Maize exudates influenced the expression of 191 genes (3.70% of the transcriptome; 43% upregulated, 57% downregulated) (Fig. 2). Moreover, changes in expression triggered by tomato exudates were characterized by a higher average in log_2_FC (2.71 log_2_FC for tiDEGs vs 2.29 log_2_FC for miDEGs) (Dataset S1). The observed difference in the total number of tiDEGs and miDEGS, as well as the difference in the average log_2_FC values between these groups, suggests that the response of P482 to the exudates of tomato was more versatile and pronounced than that to the exudates of maize.

**Fig. 2.**
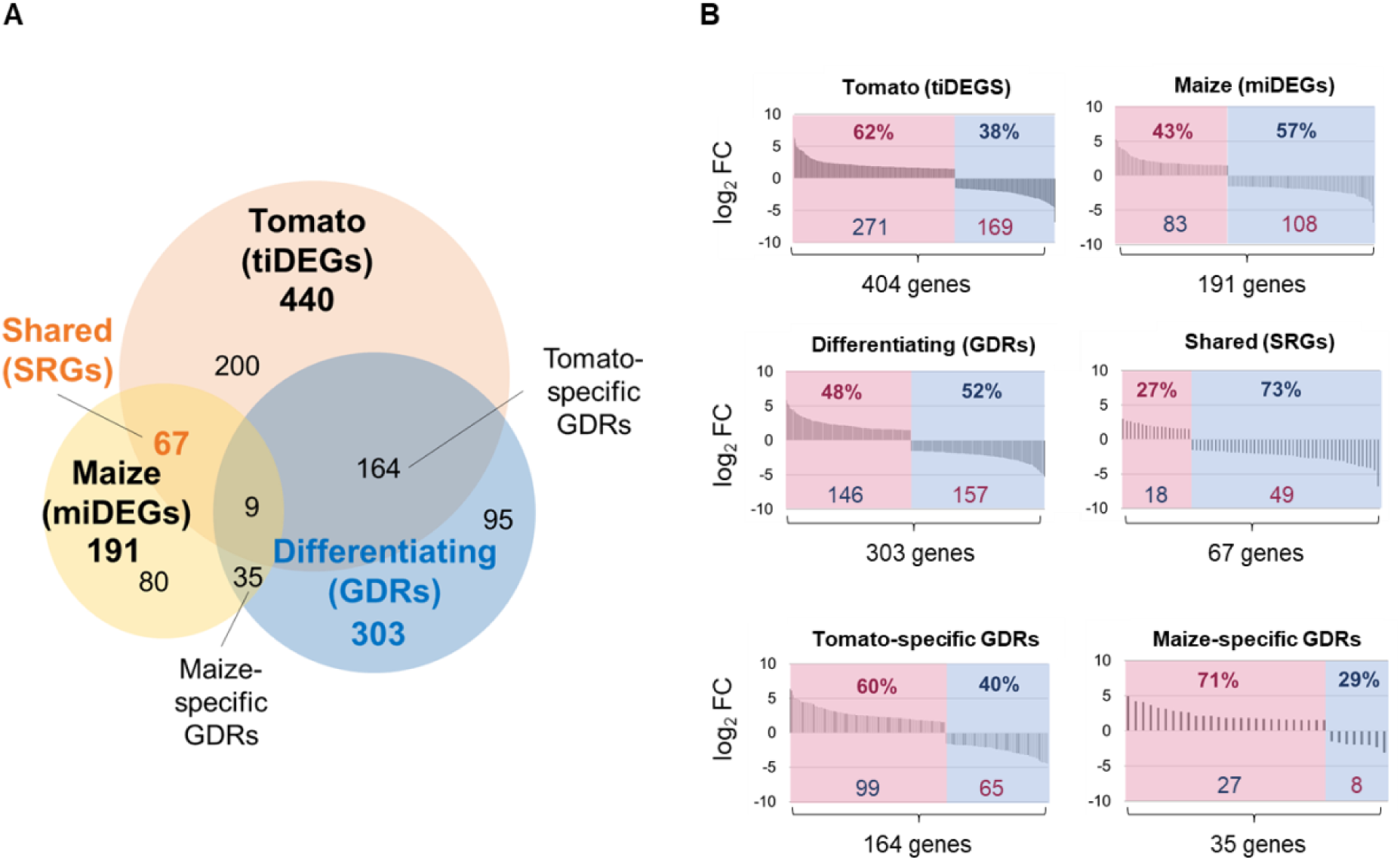
Subsets of genes differentially expressed in *P. donghuensis* P482 in response to root exudates (A) and the proportion of genes up- and downregulated in each subset (B). The shared response subset (SRGs) was established in a Venn analysis by superimpositions of three gene sets: genes with expression significantly altered in response to the root exudates of tomato (tiDEGs, 404) or maize (miDEGs, 191) and genes the expression of which was significantly different in response to the two exudate types (GDRs). The remaining subsets represent: 67 – genes of the shared response of P482 to both exudate types (SRGs), 164 – tomato specific GDRs with expression altered only by tomato exudates but not maize; 35 – maize specific GDRs with expression altered only by maize exudated but not tomato; 9 – genes with expression altered by both exudate types, yet in a different manner/to a different extent. Significance cutoffs: α=0.05; 1.5 log_2_FC. A list of loci in each subset can be found in Dataset S2. In panel B, upregulated genes are shown in magenta (on the left) and downregulated genes are in blue (on the right). Both the percentage of genes and the actual ORF count are indicated in the graphs.

Apart from identifying genes the expression of which was significantly different compared to 1C medium, we have determined genes with expression significantly different (>1.5 log_2_FC, p<0.05) between the two exudate treatments (tomato vs maize), irrespective of whether their expression was significantly different from that in 1C medium alone (untreated). Genes differently expressed in P482 when comparing the two exudate treatments were termed Genes of the Differentiating Response (GDRs). The pool of GDRs included 303 transcripts (5.86% of the transcriptome; 48% upregulated, 52% downregulated) (Fig. 2).

The three gene pools: tiDEGs (440), miDEGs (191) and GDRs (303), were superimposed, and the results were presented in the form of a Venn diagram (Fig. 2; Dataset S2, Dataset S4). The primary goal of this analysis was to identify the subset of genes of the shared response to exudates. By shared response, we understand genes the expression of which was altered in a similar manner by exudates of both plant species. The shared response included 67 genes (1.30% of the transcriptome; 27% upregulated and 73% downregulated (Fig. 2). This subset is further referred to as the Shared Response Genes (SRGs).

Superimposition of tiDEGs, miDEGs, and GDRs also revealed subsets of GDRs designated as tomato-specific GDRs (164 genes) and maize-specific GDRs (35 genes). These genes were not only among GDRs, thus showing different expression upon the two exudate treatments (>1.5 log2FC), but also their expression was significantly affected (>1.5 log2FC) upon exudate treatment when compared to 1C medium alone. Narrowing GDRs to tomato-specific and maize-specific subsets was useful to highlight the most host-specific aspects of the differentiating response. Additionally, nine GDRs were significantly altered in both tomato and maize, however differently or to a significantly different extent, excluding them from the shared response (Fig. 2). Narrowing GDRs to tomato-specific and maize-specific subsets was useful to highlight the most host-specific aspects of the differentiating response.

A complete list of genes assigned to the discussed groups, along with their annotations and recorded changes in the expression levels, is provided in Dataset S3.

### KEGG and COG enrichment analysis, ranking lists and gene networking

To identify which aspects of the metabolism of P482 were affected as part of the shared and differentiating response of P482 to exudates, we performed KEGG pathways and COG enrichment analyses for the 67 genes of the shared response (SRGs) and the 303 genes of the differentiating response (GDRs) (Dataset S5). KEGG pathways significantly enriched within SRGs included ‘Sulfur metabolism’ (8 genes) and ‘ABC transporters’ (11 genes) (Fig. 3 A), while the enriched COGs included three categories: 1) Inorganic ion transport and metabolism (18 genes); 2) Inorganic ion transport and metabolism / Signal transduction mechanism (7 genes); 3) Amino acid transport and metabolism/ Signal transduction mechanism (2 genes) (Fig. 3 B). As for GDRs, nine enriched KEGG pathways were identified: ‘Oxidative phosphorylation’ (10 genes), ‘Sulfur metabolism’ (8 genes), ‘Phenylalanine, tyrosine and tryptophan biosynthesis’ (4 genes) and ‘Thiamine metabolism’ (3 genes) and five overlapping pathways including ‘Pyruvate metabolism’ (9 genes), ‘Glyoxylate and dicarboxylate metabolism’ (9 genes), ‘Propanoate metabolism’ (14 genes), ‘Fatty acid degradation’ (5 genes), ‘Valine, leucine and isoleucine degradation’ (13 genes) (Fig. 3 C). COGs enriched within GDRs included two categories: ‘Energy production and conversion’ (35 genes) and ‘Lipid transport and metabolism’ (18 genes). A single COG category, ‘Replication, recombination and repair’, here containing only two genes (*phrB, polC*), was found to be underrepresented among GDRs (Fig 3 D).

**Fig. 3.**
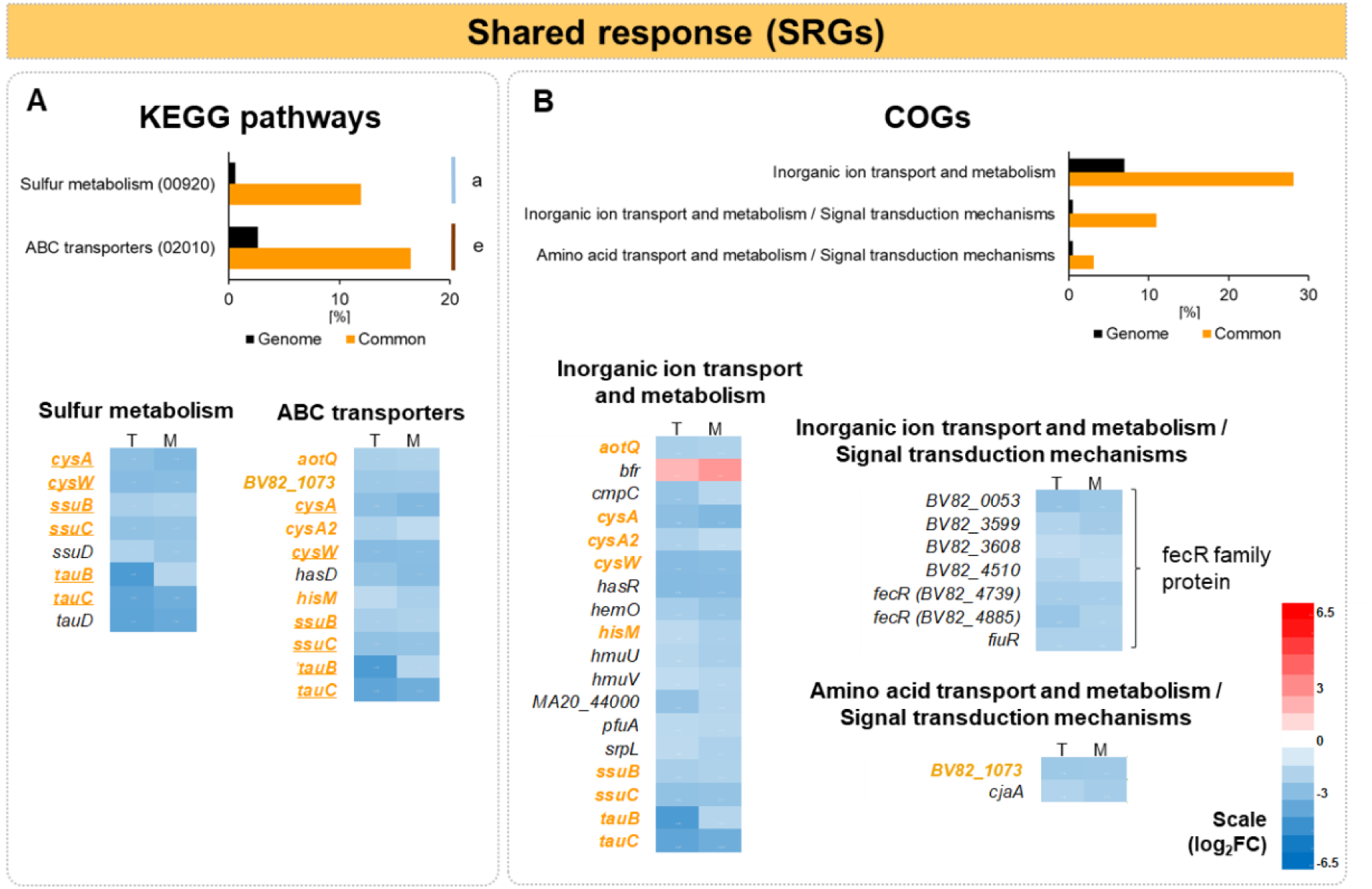
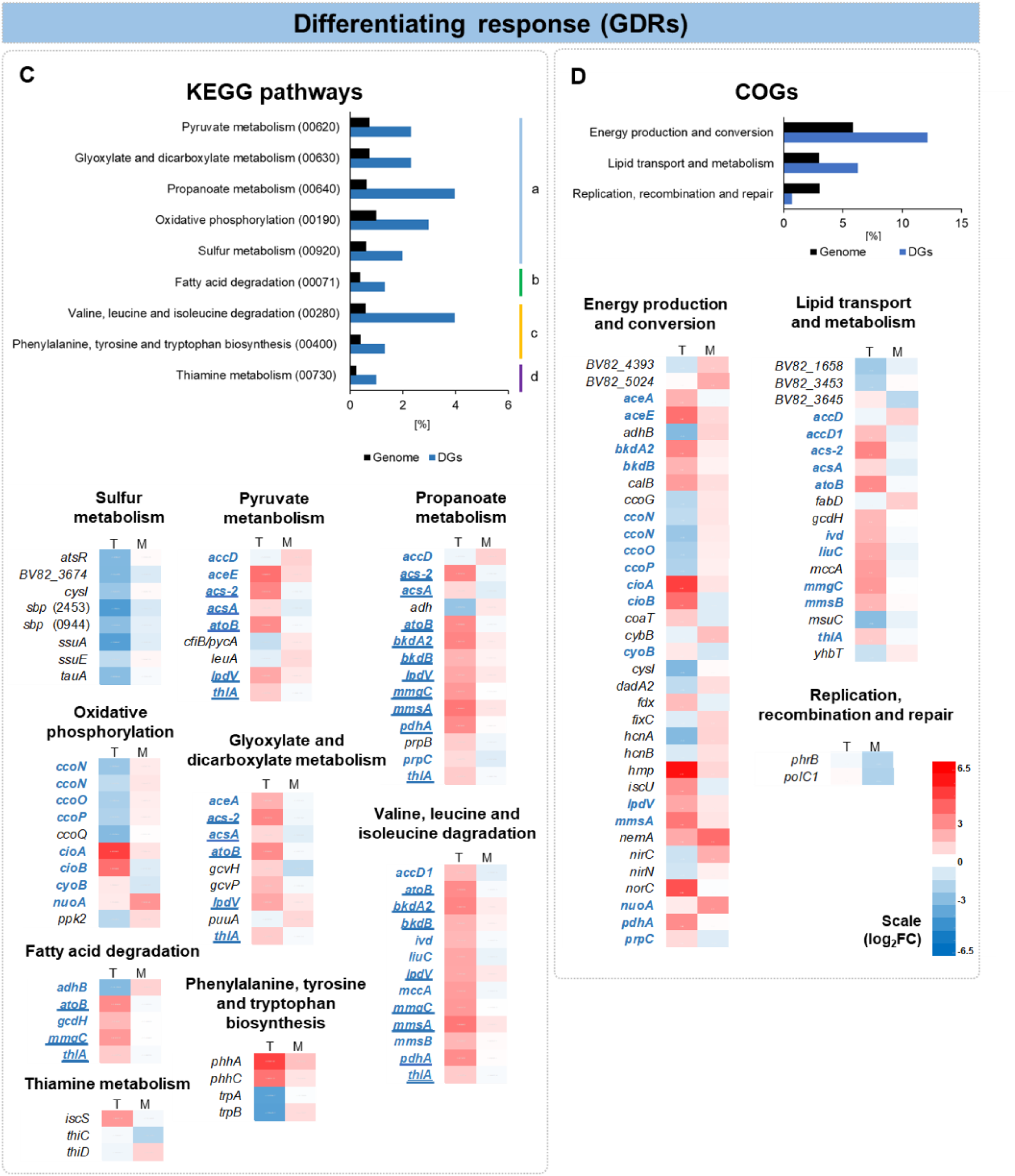
KEGG pathways and COG categories significantly enriched within the shared (A, B) and differentiating (C, D) transcriptomic responses of P482 to tomato and maize root exudates. All categories and pathways were enriched with the exception of ‘Replication, recombination and repair’ (D) which was underrepresented. For individual genes within pathways: red and blue indicate upregulation and downregulation of gene expression, respectively; the left column shows changes in expression (log_2_FC) upon addition of tomato exudates (letter T) and right column upon the addition of maize exudated (letter M). KEGG categories: a – carbohydrate metabolism, b – lipid metabolism, c – amino acid metabolism, d – metabolism of cofactors and vitamins, e – environmental information processing. Genes shown in bold color font are listed both within enriched KEGG pathways and enriched COGs. Underlined genes overlap between categories within the KEGG pathways.

An approach we applied in parallel to the enrichment analysis was to create ranking lists of genes with the highest differences in expression (up- and down-regulation) upon the exudate treatments. The reason behind it was that not all genes with high fold change in expression were a part of the enriched methabolic pathways. Moreover, the response of P482 to the exudates of tomato was more pronounced (involved more genes and with a higher average log_2_FC) than that to maize-derived compounds. As a result, the tomato-related changes in gene expression are the primary driver of variability in the pool of GDRs. To prevent a situation where maize-driven changes are underreported, we created lists of top-ranking up- and down-regulated genes not for the total pool of GDRs (where the tomato-related genes dominate) but for the tomato-specific and maize-specific GDRs (SI Appendix, Tables S5 and S6).

Another type of analysis we performed was gene networking. This approach helps to reveal associations between genes and pathways that are non-obvious or hard to track manually in a big data set. The analysis was carried out for genes within SRGs and GDRs using the STRING tool (30). Tabular presentation of the results is available in Dataset S6. Graphical representation of the networking analysis was included in a more comprehensive figure, referred to further in the text.

Below we provide the main results of our analyses, divided, for convenience, into two main sections: ‘Shared response’ and ‘Differentiating response’ and further into subsections referring to particular aspects of bacterial metabolism.

### Shared response

#### Sulfur assimilation

KEGG pathways significantly enriched within the shared response of P482 to both exudate treatments included ‘Sulfur metabolism’ (8 genes) and ‘ABC transporters’ (11 genes) (Fig. 3 A). The two pathways show a considerable overlap as five genes from the ‘ABC transporters’ (*cysA*, *cysW*, *ssuB*, *ssuC tauB*, *tauC*) encode transporters involved in the acquisition of sulfur. Noteworthy, all genes in the KEGG pathways enriched within SRGs were significantly downregulated in the presence of both types of exudates compared to the growth in the unsupplemented medium (Table 1)

Products of genes representing the enriched pathways include CysW and CysA, a membrane protein and an ATPase, respectively, which are the components of the sulfate-thiosulfate permease (SulT). Sulphate and thiosulphate represent inorganic sources of sulphur, preferentially used by bacteria (35). Located next to the *cysA* (BV82_2456) and *cysW* (BV82_2455) was another downregulated gene, BV82_2457, represening the enriched COG category ‘Inorganic ion transport and metabolism’. BV82_2457 encodes a type I phosphodiesterase / nucleotide pyrophosphatase family protein. The catalytic core of nucleotide pyrophosphatases/phosphodiesterases is known to be present in some arylsulphatases (36).

Other genes of the enriched sulfur-related pathways included *tauBC* and *ssuBC* encoding TauBC and SsuBC. These proteins are involved in the transport of taurine and other alkanesulfonates, respectively. Alkanesulfonates are organic compounds that can serve as alternative sources of sulfur for bacterial cells in the absence of inorganic sulfate or cysteine (35). Apart from the abovementioned transporter genes, sulfur metabolism pathway downregulated in P482 as part of the shared response also involved *ssuD* and *tauD*, the products of which are involved in the release of sulfur from the sulfur-containing compounds following their import into the cell (37). The *tauB*, *tauD* and *tauC* genes, apart from being part of the enriched KEGGs and COGs, were also among GDRs with the most substantial shift towards downregulation.

Gene networking analysis performed by STRING revealed the connection between *cys*, *ssu* and *tau* genes (Fig. 4 A). Additionally, STRING enabled the identification of another downregulated gene presumably involved in sulfur acquisition in P482, the *sfnR*. In *Pseudomonas putida* DS1, the *sfnR* gene was found to be necessary for the utilization of dimethyl sulfide (DMS) as a sulfur source (38).

**Fig. 4.**
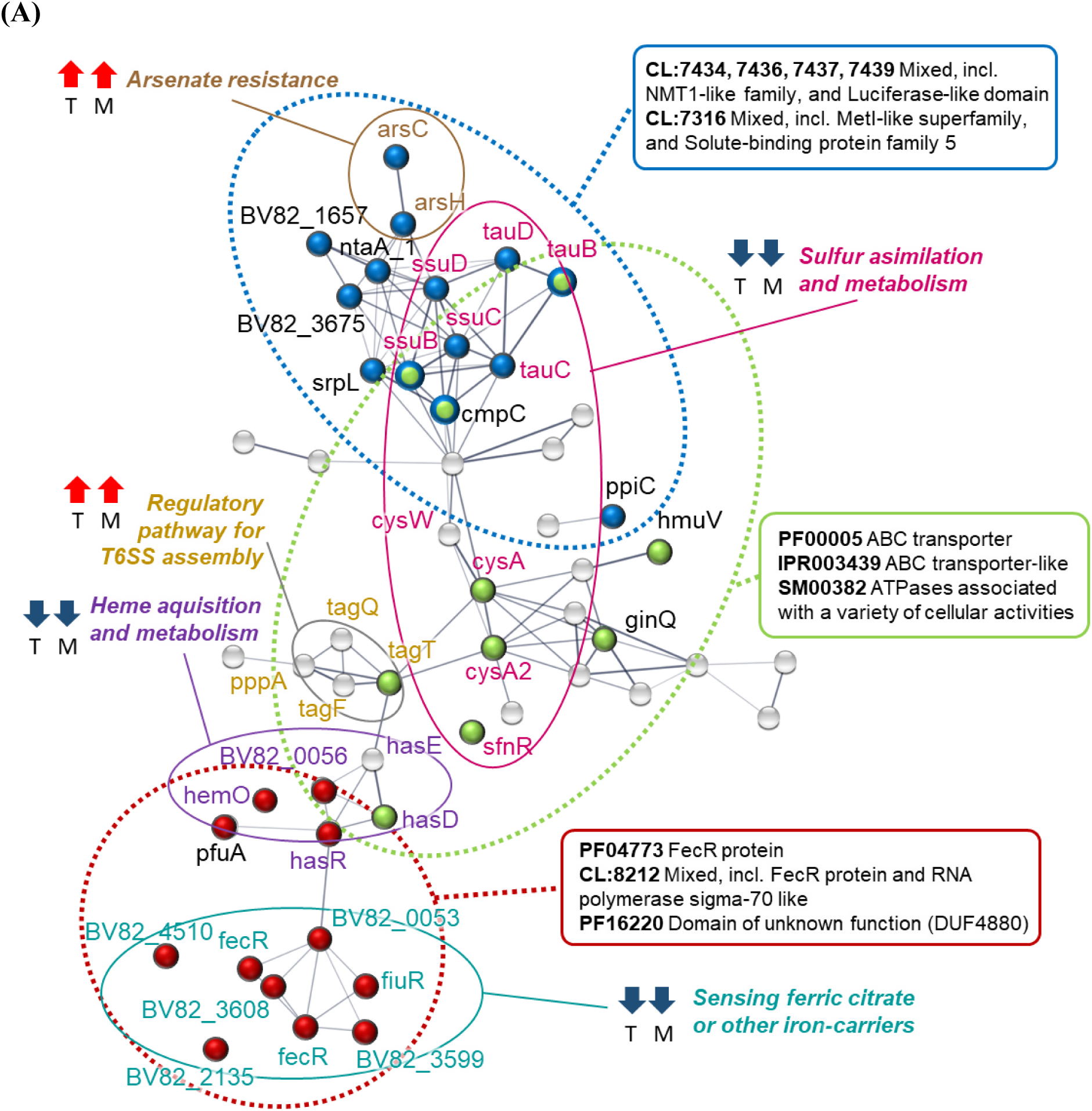
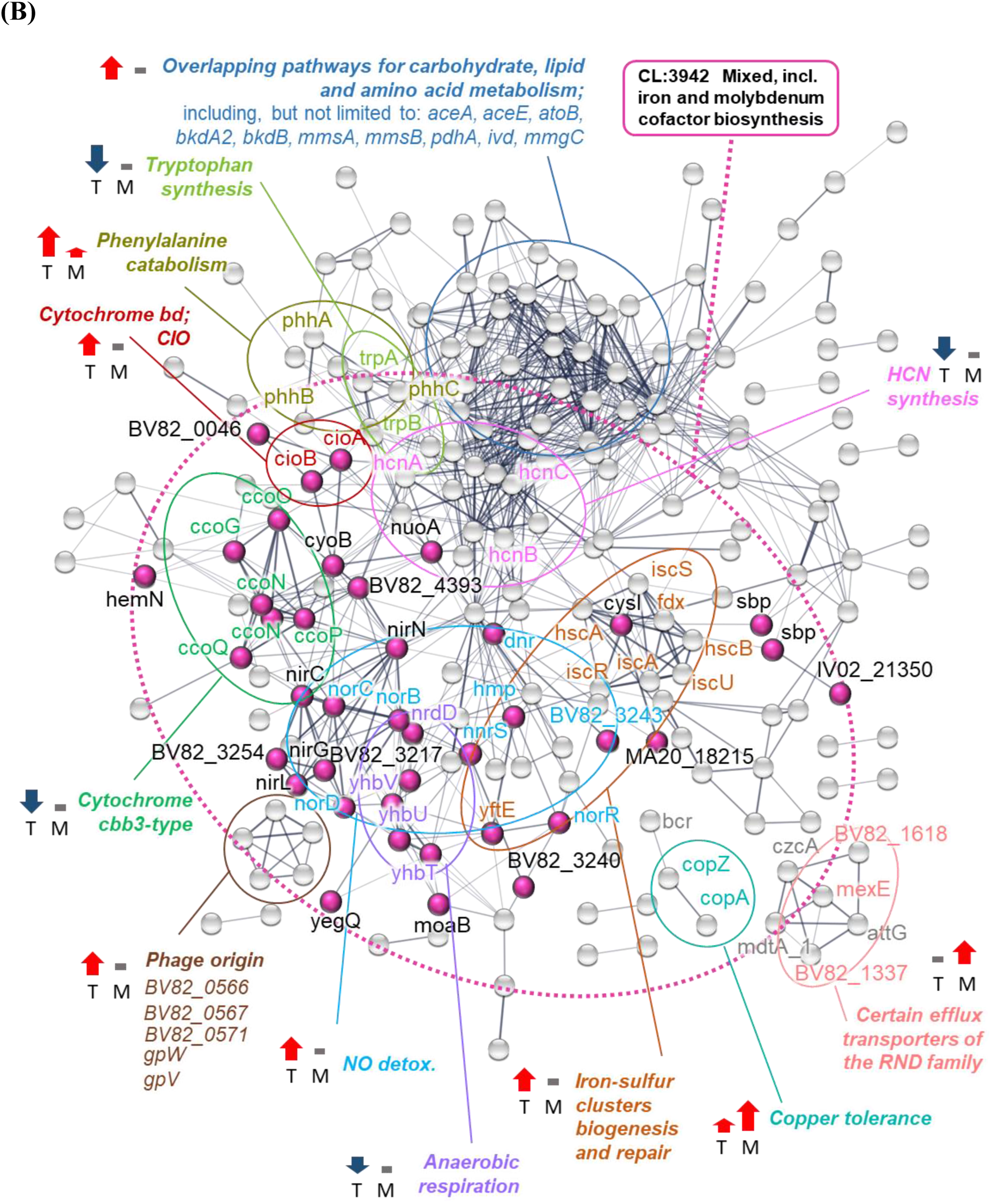
Gene networking for the 67 genes of the shared (A) and 303 genes of the differentiating (B) transcriptomic response of P482 to plant root exudates. The analysis was performed using STRING and the image was manually curated. Each node in the graph represents a gene of P482 with a name assigned with eggNOG. Filled-in nodes comprise enriched clusters identified by STRING. Designations of the enriched clusters are indicated in frames colored to match the nodes, with dotted line surroundings added for extra guidance. Based on literature, we hypothesize that certain groups of genes are involved in particular metabolic processes. Names of these genes are written in color corresponding to that of the name of the process. Red arrow up indicates genes upregulation, blue arrow indicates genes downregulation, dash indicates no significant change of expression from the medium alone; this is shown for the growth of P482 in both tomato (T) and maize (M) exudates. Network edges indicate confidence. Interaction based on all available sources for version 11.5. Minimal interaction score 0.4 (medium). Disconnected nodes were unabled, with the exception of genes that were found to be a part of enriched clusters. Each gene can be matched with a loci designation using Dataset S1.

Taken together, the obtained results show that, in our experimental setup, both types of exudates decrease the expression of multiple genes related to the acquisition of sulfur.

#### Arsenic resistance

Among genes with the highest upregulation among the shared response genes were the *arsC* and *arsH* (between 2.35-2.66 log_2_FC) (SI Appendix, Table S7). Both genes are related to the metabolism of arsenic – a highly toxic metalloid ubiquitous in the environment (39). ArsC protein is an arsenate reductase transforming arsenate (As(V)) to arsenite As(III) in a first step of arsenic detoxification. It was shown in *Eschericha coli* that the resulting As(III) is extruded from the cells by the ArsA/ArsB, an ATP-driven pump (40). Homologues of *arsA* and *arsB* were not found among the SRGs of P482. The second arsenic-related gene upregulated in P482 encodes ArsH, one of the more recently described Ars proteins. It is an organoarsenical oxidase enzyme that was found to confer resistance to trivalent forms of the herbicide monosodium methylarsenate due to the oxidation of the trivalent species to pentavalent species (41). Within the shared response of P482 to exudates, gene networking analysis revealed a link between the sulfur assimilation genes and the arsenic resistance genes *arsC* and *arsH* (Fig. 4 A).

#### Iron acquisition and intercellular iron homeostasis

##### Sensing of xenosiderophores

One of the COG categories enriched within the shared response was a double-function category “Inorganic ion transport and metabolism / Signal transduction mechanism”. In this study, this category was represented solely by seven *fecR*-like genes (BV82_0053, BV82_3599, BV82_3608, BV82_4510, BV82_4739, BV82_4885, BV82_ 1854 (*fiuR*)). FecR proteins are transmembrane sensors involved in signal transduction. In *Escherichia coli*, FecR protein cooperates with extracytoplasmic-function sigma factor FecI and receptor protein FecA to direct the expression of the *fecABCDE* operon involved in the uptake of ferric citrate (42). However, bacterial species can harbour many *fecR*-like genes. In the genome of *P. aeruginosa*, there are fourteen *fecR*-like genes located adjacent to iron-starvation sigma factors that can potentially be induced in the presence of cognate iron carriers (43). The observed downregulation of *fecR*-like genes in *P. donghuensis* P482 following growth in the presence of root exudates could imply either a lower availability of the cognate compounds and/or, more probably, lower demand for the import of xenosiderophores. In relation to the transport of iron carriers into the cell, we also observed downregulation of *pfuA* (BV82_0628) gene encoding a TonB-dependent siderophore receptor family protein.

To further address the matter of how the addition of exudated influences iron availability to P482 in our experimental setup, we have investigated the expression levels of 3 genes: BV82_1009, an NRPS-type synthase involved in the synthesis of a potent siderophore pyoverdine; BV82_1008 (*pvdS*) – a sigma factor involved in the regulation of synthesis of pyoverdine, and BV82_4709 involved in the synthesis of 7-hydroxytropolone with iron-scavenging properties. In all three cases, the expression of the investigated genes in the presence of both types of exudates did not significantly differ (log2FC <1.5) from the expression in 1C medium alone (Dataset S1). However, in our recent study (44) we have found that the expression of BV82_1009 and BV82_4709 can be detected in unsupplemented 1C medium, here used as the reference (background), therefore the results from the RNAseq experiment can only be interpreted in terms the influence of exudates on the existing expression status of these genes, and they do not provide the information wheather the basal expression is high or low.

##### Heme acquisition and metabolism

Other genes assigned to COG ‘Inorganic ion transport and metabolism’ included genes involved in the uptake or metabolism of heme (*hasR*, *hemO*, *hmuU* and *hmuV*).

Among the top-ranking downregulated SRGs, the gene showing the most significant downregulation was BV82_0056 encoding a heme-binding A family protein (−6.8 log_2_FC) (SI Appendix, Table S7). Downregulated genes involved in the uptake or metabolism of heme were also represented within the enriched COG category ‘Inorganic ion transport and metabolism’ (*hasR*, *hemO*, *hmuU* and *hmuV*) and the ‘ABC transporter’ pathway (*hasD*). HasR is a member of the TonB-dependent family of receptors which can transport either heme itself or heme bound to HasA (45). HemO is a heme oxygenase that can perform oxidative cleavage of heme and enable some bacteria to acquire heme-derived iron for further use (46). HmuUV is an ABC-type transporter that allows the import of heme bound to its cognate substrate-binding protein HmuT (47). The product of the last enlisted gene, HasD, is a part of ATP-binding cassette exporter HasDEF for the heme acquisition protein HasA.

The STRING networking analysis revealed a link between the genes of heme acquisition and metabolism (BV82_0056, *hemO*, *hasD*, *hasE*, *hasR*) and the *fecR* genes (BV82_4739, BV82_4885, BV82_4510, BV82_1854, BV82_0053, BV82_3608, BV82_3599) involved in the sensing ferric citrate and potentially other usable iron-carriers. Furthermore, the expression of all of these genes related to the acquisition of different forms of iron was downregulated in the presence of root exudates of both plants compared to growth in an unsupplemented medium.

##### Intracellular iron homeostasis

While the expression of *fecR* genes for the detection of xenosiderophores and the heme acquisition genes was downregulated as part of the shared response of P482 to exudates, the expression of *bfr* gene encoding bacterioferritin was upregulated (1.87 and 2.67 log_2_FC for tomato and maize, respectively). Bacterioferritin is a cytosolic iron storage protein shown to have a crucial role in maintaining iron homeostasis in *Pseudomonas aeruginosa* (48, 49).

#### Assembly of T6SS

Upon stimulation with both types of exudates, we observed upregulation of expression of *tagT, tagQ,* BV82_1738 (*tagF*) and *pppA* genes. Based on STRING networking, these genes are linked to each other and the *fecR*-heme node. In *P. aeruginosa*, TagT, TagQ and TagF are proteins involved in the regulatory cascade that controls the activation of the type VI secretion system (T6SS). This secretion system enables bacteria to inject proteins into other bacterial cells and is best known for its role in interbacterial competition (50, 51). In *P. aeruginosa,* protein secretion is regulated post-translationally by phosphorylation of threonine, with PpkA kinase required for the assembly of T6SS. The activity of the kinase antagonized by the PppA phosphatase (52). In the shared response of *P. donghuensis* P482 to root exudates, we observed increased expression of *pppA* and *tagF.* Both of these genes are considered negative regulators of T6SS (53). Therefore, results obtained for P482 suggests overall downregulation of T6SS. However, the regulation of T6SS assembly in different bacterial species remains poorly understood (54).

#### Transport of amino acids and the shikimate pathway

Genes within the enriched KEGG ‘ABC transporter’ pathway unrelated to sulfur metabolism included *aotQ*, *hisM* and a nameless gene in locus BV82_1073, all of which were downregulated upon exudate treatments (−2.18, −1.69 and −2.49, respectively). Predicted products of these genes are AotQ, an arginine/ornithine transport protein involved in the movement of these nitrogen-containing compounds, and HisM, a part of HisQMP_2_ transporter for positively charged amino acids (lysine, arginine, histidine), first described for *Salmonella enterica* serovar Typhimurium (55). The last gene, BV82_1073, encodes a bacterial solute binding protein, presumably involved in amino acid transport and metabolism and/or signal transduction mechanisms. Genes *aotQ*, *hisM* and BV82_1073 were also present among the enriched COGs – the first two in the ‘Inorganic ion transport and metabolism’ and BV82_1073 in the ‘Amino acid transport and metabolism/ Signal transduction mechanism’ category. In the latter, the BV82_1073 was accompanied by another downregulated gene, BV82_2469, the product of which is annotated as transporter substrate-binding domain-containing protein with 45% identity (88% query coverage) to a highly immunogenic, amino acid-binding protein CjaA from *Campylobacter jejuni* (56). Although it is challenging to speculate on the overall meaning of the abovementioned changes in gene expression, it is clear that the addition of both types of exudates led to the downregulation of certain amino acid transport paths in strain P482.

Additionally, concerning amino acids, one of the top-ranking downregulated SRGs was the *aroF* (BV82_3002; −3.10 and −3.35 log_2_FC in response to tomato and maize, respectively), the product of which, 3-deoxy-7-phosphoheptulonate synthase (DAHP), is the first enzyme of the shikimate pathway responsible, for the synthesis of folates (vitamin B_9_) and aromatic amino acids: phenylalanine, tyrosine and tryptophan, among other metabolites.

### Differentiating response

#### Nitric oxide detoxification and the repair of iron-sulfur clusters

One of two genes that showed the most substantial upregulation among the genes of the differentiating response (GDRs) was the *hmp* (BV82_4743) encoding flavohemoglobin (Hmp). The expression of this gene was significantly increased (6.13 log_2_FC) in *P. donghuensis* P482 grown in the presence of tomato exudates but not those of maize origin (1.05 log_2_FC). Hmp is a well-documented component of bacterial response to prevent the toxic effects of nitric oxide (NO). Under aerobic conditions, the Hmp enzyme catalyzes the reaction of NO with oxygen to yield nitrate (NO_3_-) (57).

A similar pattern of changes in gene expression as the one observed for the *hmp* was also true for *norD*, *norC* and *norB* genes encoding the nitric oxide reductase (NOR). This membrane-integrated enzyme catalyzes the reduction of nitric oxide (NO) to nitrous oxide (N_2_O) (58). NOR plays a vital role in nitrogen metabolism in denitrifying bacteria, as well as it constitutes another, next to Hmp, important NO-detoxifying mechanism (59).

In denitrifying bacteria, NO is formed as an intermediate in a stepwise reduction of nitrate, through nitrite, to nitrogen oxide. In the course of denitrification, electrons from the respiratory chain are transferred to a non-oxygen acceptor, making denitrification a bacterial strategy to generate energy under conditions of limited oxygen availability (60). Among those *Pseudomonas* spp. which can perform denitrification, such as *P. aeruginosa* and *P. stutzeri*, the formation of NO from nitrite in the second step of the process requires the activity of NirS (61, 62). The NirS protein is a periplasmic cytochrome cd_1_ nitrite reductase carrying heme c and heme d_1_ cofactors (63). *P. donghuensis* P482 does possess a homologue of *nirS* (BV82_3250) and homologues of multiple other genes of the *nir* cluster (*nirC, nirF, nirL, nirG, nirH, nirN, nirB, nirD, nirJ, nirM*). However, the expression of *nirS* was not significantly altered in the presence of maize exudates nor the presence of exudates of tomato. The expression of other *nir* genes was also not significantly upregulated in the presence of exudates, independent of their plant origin. This implies that the strong upregulation of activity of NOR, observed in P482 in the presence of tomato exudates, is not related to an increased indigenous production of NO due to a switch of the strain to a denitrification pathway.

Genes encoding two NO-responsive transcription factors, NorR and Dnr (BV82_3238 and BV82_3242), are located adjacent to the *nor* genes in *P. donguensis* P482. Both of these proteins were reported to activate the transcription of NOR, with Dnr (dissimilative nitrate respiration regulator) also reported to modulate the expression of other denitrification-related genes *nirS* and *nosZ* (61, 64). In P482, in the presence of tomato exudates, we observed a significant upregulation of expression of *dnr* (3.23 log_2_FC) while the expression of *norR* is downregulated (−1.67 log_2_FC). In the presence of maize exudates, the expression *dnr* is not affected when compared to the 1C medium alone, while in the case of *norR* we noticed a tendency towards upregulation (1.41 log_2_FC), although below the adopted significance threshold (>1.5 log_2_FC). Although NorR and Dnr are definitely regulatory proteins involved in the management of nitrosative stress, it seems to depend on bacterial species which regulons these proteins control and wheather this regulation is connected to response to environmental factors other than nitrosatite stress, including oxygen availability and iron metabolism (65, 66).

Encoded by locus BV82_3241, therefore located between *norR* and *dnr* regulatory genes (BV82_3238 and BV82_3242, respectively), is the *nnrS* gene. NnrS is a heme- and copper-containing transmembrane protein that contributes to NO stress tolerance in *Vibrio cholerae* (67). In P482, the *nnrS* was significantly upregulated (4.99 log_2_FC) in the presence of tomato exudates.

Another protein with a well-established link to bacterial resistance to nitrosative stress is the YtfE (66). Although this protein is not directly involved in the turnover or sensing of NO, it does play an important role in the metabolism/repair of iron-sulfur (Fe-S) clusters (68). Homologues of *ytfE* can be found in numerous bacteria (69), and the gene was shown to be consistently and massively upregulated upon bacterial exposure to nitric oxide (66). In P482, the *ytfE* gene shows the highest upregulation among GDRs, being upregulated by 6.36 log_2_FC in the presence of tomato-derived compounds. Proof of a strong link between *yftE* and other genes related to nitrosative stress in strain P482 is that *yftE* (BV82_3239) is encoded next to *norR*, *nnrS*, *dnr*, *norD*, *norB* and *norC* in a single cluster spanning from BV82_3238 to BV82_3246. Intriguingly, the cluster also contains two strongly upregulated genes encoding proteins with only a vague annotation: “putative membrane protein” and “putative DnrP protein” (BV82_3240 and BV82_3243, respectively).

Moreover, in P482 grown in the presence of tomato exudates, we observed not only the upregulation of *yftE* involved in the repair of Fe-S clusters but also upregulation of genes encoding proteins related to sulfur cluster biogenesis: *iscR*, *iscS*, *iscU*, *iscA*, *hscB*, *hscA, fdx*(70). In P482, these genes are encoded by loci BV82_0252 to BV82_0258.

#### Terminal oxidases and respiration

One of the KEGG pathways enriched within GDRs was ‘Oxidative phosphorylation’. Pseudomonads harbor several terminal oxidases that can be used alternatively to reduce oxygen to water during aerobic respiration (71, 72). This allows *Pseudomonas* bacteria to adapt their respiratory chain to best suit given conditions. In response to tomato exudates, strain P482 showed a significant downregulation of *ccoN*, *ccoO*, *ccoQ* and *ccoP* genes encoding the subunits of a *cbb3*-type cytochrome c oxidase. This effect was not observed for the exudates of maize.

While *ccoN*, *ccoO*, *ccoQ* and *ccoP* genes were downregulated in P482 in the presence of tomato exudates, the *cioA* and *cioB* genes were significantly upregulated (4.9 and 3.7 log_2_FC, respectively). The CIO stands for <u>c</u>yanide-insensitive <u>o</u>xidase and the *cioA* and *cioB* genes encode the two subunits of cytochrome *bd* complex. Unlike in heme-copper oxidases, the active centre of the quinol CIO oxidases consists solely of heme (72). This type of oxidases is known to be harboured by some cyanide-tolerant bacteria (73). Cyanide is a toxic compound that rapidly blocks respiration mediated by heme-copper cytochrome c oxidases such as the *cbb3*-type cytochrome c oxidase. However, microorganisms which can switch to the alternative CIO pathway can withstand high cyanide concentrations or even use it for growth as a sole nitrogen source (72). Strain P482 was shown to produce hydrogen cyanide (15). However, in P482 exposed to tomato exudates, genes *hcnA* and *hcnB* encoding the hydrogen cyanide synthase were downregulated, implying that upregulated expression of cyanide-insensitive cytochrome *bd* in this strain does not occur to counter the toxicity of high levels of self-produced hydrogen cyanide.

The enriched ‘oxidative phosphorylation’ subset in the GDRs of P482 contained almost exclusively genes with expression significantly affected in the presence of tomato-derived compounds. However, in the presence of maize exudates, we observed upregulation of *nuoA* (2.77 vs 0.54 log_2_FC, respectively). The *nuoA* encodes one of the subunits of the NADH dehydrogenase of complex I that shuttles electrons from NADH, via FMN and iron-sulfur centers, to quinones in the respiratory chain (74) (https://www.uniprot.org/uniprot/Q02NC9).

#### Catabolism of amino acids/fatty acids, conversion of alpha-keto acids to CO_2_ and the glyoxylate shunt

Five enriched KEGG pathways within GDRs showed a considerable overlap. This included ‘Pyruvate metabolism’ (9 genes), ‘Glyoxylate and dicarboxylate metabolism’ (9 genes), ‘Propanoate metabolism’ (14 genes), ‘Fatty acid degradation’ (5 genes) and ‘Valine, leucine and isoleucine degradation’ (13 genes). Although the sum of genes in the listed pathways equals 50, there are only 28 unique genes in this group, several appearing in more than a single pathway. These genes also appear within the enriched COGs assigned to either ‘Energy production and conversion’ or ‘Lipid transport and metabolism’ (Fig. 3 C, D). The expression of majority of genes in the described group was altered in the presence of the exudates of tomato.

Tomato-derived compounds upregulated the expression of four cluster-forming genes in loci BV82_0838-0841: *lpdV* for dihydrolipoyl dehydrogenase (2.26 log_2_FC), *bkdB* (1.95 log_2_FC) for lipoamide acyltransferase component of branched-chain alpha-keto acid dehydrogenase complex and *bkdA2* (3.14 log_2_FC) and *pdhA* (2.94 log_2_FC) encoding beta and alpha subunits of 2-oxoisovalerate dehydrogenase, respectively. Protein products of these genes are the components of branched-chain alpha-keto dehydrogenase complex – one of three multienzyme complexes metabolizing pyruvate, 2-oxoglutarate, and branched-chain 2-oxo acids. The size of these multienzyme complexes ranges from 4 and 10 million Da, placing them among the largest and most sophisticated protein assemblies known (75). The three complexes show structural and functional similarities. They all comprise E1, E2 and E3 components, each present in numerous copies and utilizing multiple cofactors (75). The overall function of these complexes is the conversion of alpha-keto acids to acyl-CoA and CO_2_. The important representatives of alpha-keto acids include pyruvate, a pervasive metabolic intermediate, oxaloacetate, a component of the Krebs cycle, and alpha-ketoglutarate. Alpha-keto acids often arise by oxidative deamination of amino acids, and, reciprocally, they are the precursors for the synthesis of thereof. The 2-oxoisovalerate dehydrogenase, which in P482 is encoded by the abovementioned *bkdA2* and *pdhA* genes, is an enzyme known to participate in the degradation of valine, leucine and isoleucine (76). In parallel to upregulation of *bkdA2* and *pdhA*, we see upregulation of *mmsA* (3.49 log_2_FC) and *mmsB* (1.87 log_2_FC). These genes encode methylmalonate-semialdehyde dehydrogenase and 3-hydroxyisobutyrate dehydrogenase, respectively. In *P. aeruginosa* PAO, the *mmsAB* operon was shown to be involved in the metabolism of valine and mutation of these genes led to very slow growth of the strain on valine/isoleucine medium (77).

Moreover, in the presence of tomato exudates, we saw downregulation of BV82_3254. Although this particular GDR was not among genes of the enriched pathways, it is interesting as it encodes a putative Lrp/AsnC family transcriptional regulator. This family of regulators refers to proteins orthologous or paralogous to the *E. coli* leucine-responsive regulatory protein (Lrp) and asparagine synthase C gene product (AsnC) – two of the *E. coli* feast/famine regulatory proteins (FFRPs) (78). In *E. coli*, Lrp either represses or activates the transcription of numerous genes, in many cases depending on the extracellular availability of leucine. A high concentration of leucine in the environment is considered a proxy for sensing a nutrient-rich environment. In these conditions of ‘feast’, as opposed to ‘famine’*, E. coli* represses pathways enabling autotrophic synthesis of nutrients and shifts to a more heterotrophic mode by activating their absorption from the environment, accompanied by altered infectivity and the formation of pili (78). In P482, the expression of BV82_3254 (*lrp*) differed upon the two exudate treatments, with a tendency for downregulation in the presence of tomato exudates and up-regulation in the presence of maize exudates (Table 1).

Among pathways enriched within the differentiationg response we also found gene clusters BV82_1123-1126 and BV82_2129-2132. Genes in both clusters are upregulated in the presence of tomato exudates. The BV82_1123-1126 encodes an AMP-binding acyl-CoA ligase (here designated Acs-2), acetyl-CoA C-acetyltransferase AtoB which catalyzes the final step of fatty acid oxidation in which acetyl-CoA is released and the CoA ester of a fatty acid two carbons shorter is formed, and acyl-CoA dehydrogenase MmgC. The second cluster, BV82_2129-2132, encodes acyl-CoA dehydrogenase Ivd, carboxyl transferase AccD1, enoyl-CoA hydratase/isomerase LiuC and a HlyD secretion protein MccA. Both of these clusters contain genes present in more than one or, like in the case of *atoB*, in all five enriched KEGG pathways discussed in this paragraph. Noteworthy, both clusters encode acyl-CoA dehydrogenases. The acyl-CoA dehydrogenases catalyze the α,β-dehydrogenation of acyl-CoA esters during the catabolism of fatty acids and amino acids (79).

In response to tomato-derived compounds strain P482 showed upregulated expression of *aceA* gene encoding isocitrate lyase (2.00 log_2_FC). Upregulation of *aceA* indicates activation of the glyoxylate shunt (GS) – a two-step anaplerotic pathway (isocitrate lyase, AceA; and malate synthase, GlcB). The GS bypasses the carbon dioxide-producing steps of tricarboxylic acid cycle (TCA) and diverts part of the carbon flux at isocitrate. Complete oxidation of acetyl-CoA to CO_2_ in TCA requires the formation of citrate from the condensation of acetyl-CoA with oxaloacetate. This is because, in each turn of the TCA cycle, two molecules of CO_2_ are lost and, for the cycle to continue, they have to be replenished, and citrate needs to be recycled back to oxaloacetate. The classical TCA cycle cannot assimilate carbon and recycle citrate when acetyl-CoA is the only available carbon source. Consequently, when acetate or fatty acids are the primary sources of carbon and energy available to the cell, many bacteria, including *Pseudomonas* spp., activate an alternative pathway – the glyoxylate shunt (reviewed in: (80) and (81).

In the presence of tomato exudates we also saw upregulation of *aceE/mdeB* (BV82_4665; 3.65 log_2_FC) and *mdeA* (BV82_4666; 4.47 log_2_FC). AceE of the E1 component of the pyruvate dehydrogenase complex. In *P. putida*, it was shown that MdeB is a homologue of AceE yet with specificity for alpha-ketobutyrate rather than pyruvate (82). The MdeB may play an important role in the metabolism of alpha-ketobutyrate produced by MdeA, a methionine gamma-lyase, in strain P482 encoded by the adjacent gene highly upregulated by tomato exudates (4.47 log_2_FC). However, the MdeA can be multicatalytic and perform other reactions on other substrates, that is γ-replacement of L-methionine with thiol compounds, α, γ-elimination and γ-replacement reactions of L-homocysteine and its S-substituted derivatives, O-substituted-L-homoserines and DL-selenomethionine, and, to a lesser extent, α,β-elimination and β-replacement reactions of L-cysteine, S-methyl-L-cysteine, and O-acetyl-L-serine (https://www.uniprot.org/uniprot/P13254).

Tomato exudates also induced the catabolism of other amino acids. Genes involved in the catabolism of phenylalanine, *phhA*, *phhB* and *phhC* were significantly upregulated (4.82, 4.36 and 3.58 log_2_FC, respectively), suggesting increased degradation of phenylalanine. Addidianally, the expression *gcvH* and *gcvP* genes of the glycine cleavage system was higher in response to tomato-derived compounds than in response to maize.

The majority of genes affected by tomato exudates were upregulated. This, however, was not true for *adh* (BV82_5184) and *adhB* (BV82_5069) which, were downregulated (−2.68 and −3.12 log_2_FC) upon tomato exudate treatment. Adh and AdhB are L-threonine dehydrogenase, a zinc-binding protein, and an iron-containing alcohol dehydrogenase family protein, respectively. Downregulation of *adh* and *adhB* can indicate a tendency for decreased activity of at least some alcohol dehydrogenases (ADHs) in P482 in the presence of tomato-derived compounds.

To conclude, the pathways activated in P482 in the presence of tomato exudates imply that in this treatment, the primary carbon sources available to the strain at the point of cell harvest were compounds yielding acyl-CoA, most probably amino acids and/or fatty acids.

#### Amino acid synthesis: tryptophan and methionine

Tomato exudates significantly influenced pathways for the synthesis of two amino acids: tryptophan and methionine. Tryptophan is a proteinogenic aromatic amino acid which can also be a precursor of signalling molecules such as the plant hormone indole-3-acetic acid (IAA), kynurenine and the *Pseudomonas quinolone signal* (PQS) (83). In the presence of tomato exudates, the expression of genes encoding tryptophan synthase, *trpA* and *trpB*, was significantly downregulated (−4.21 and −4.28 log_2_FC, respectively), implying reduced synthesis of tryptophan by P482. Contrary, we observed significant upregulation of *metE* (BV82_4439; 4.73 log_2_FC). MetE is a 5-methyltetrahydropteroyltriglutamate—homocysteine S-methyltransferase that catalyzes the transfer of a methyl group from 5-methyltetrahydrofolate to homocysteine resulting in the formation of methionine.

#### Substrate binding proteins for sulfur-containing compounds

In this study, we showed that downregulation of the sulfur assimilation pathways is typical for the shared response of P482 to both tested exudates. However, a fraction of sulfur-related genes were found among GDRs. This concerns genes encoding substrate-binding proteins (SBPs) of several sulfur assimilation pathways, including *spb-*like genes BV82_2453 and BV82_0944 for obtaining sulfur from inorganic sources, *tauA* and *ssuA* – genes encoding periplasmic binding proteins for taurine and other alkanesulfonates, respectively, the BV82_3674 gene presumably encoding a substrate-binding protein aliphatic sulfonates, and the *atsR* gene reported to encode a putative periplasmic sulfate ester-binding protein in *P. putida*(84). The expression of all these genes was significantly downregulated in response to tomato but not maize. Moreover, gene BV82_3674 was the most downregulated tomato-specific GDR (−4.48 log_2_FC) (SI Appendix, Table S5). Other genes related to the sulfur metabolism that were among GDRs and not, alike most sulfur-related genes, among SRGs, include *cysI* and *ssuE* encoding the NADPH-dependant sulfite reductase and the NAD(P)H:FMN oxidoreductase, respectively. The *cysI* and *ssuE*, similarly to the substrate-binding proteins, are down-regulated only during growth in the presence of tomato exudates.

To conclude, the obtained results show that the sulfur metabolism pathways are downregulated in response to both investigated types of exudates. However, the sulfur demand of P482 seems to be reduced even further in the presence of tomato exudates.

#### Genes of phage origin

In the presence of tomato exudates but not maize, we observed a significant upregulation of genes BV82_0566, BV82_0567, BV82_0568 (gpV) BV82_0569 (*gpW*), BV82_0571. Based on annotation, these genes encode a putative holin, a hypothetical protein, a phage baseplate assembly V family protein, a lysozyme family protein and a phage tail protein I, respectively. The listed genes are definitely of phage origin, although they are just remnants of a phage that was once incorporated into the genome of the strain. Our recent *in silico* screening of a complete genome of P482 for the presence of prophages did not reveal an intact/complete prophage in this particular region of the strain’s genome (85). Noteworthy, the fact that genes of phage origin are induced in the presence of tomato exudates is another premise that the cells of P482 experience some form of stress. Prophage induction is known to occur following treatment with toxic chemicals or UV light (86).

#### Efflux pumps of the RND family

Five genes genes encoding Membrane Fusion Protein (MFP) subunits of efflux transporters from the Resistance-Nodulation-Division (RND) family showed different expression in response to the two tested exudate types. Those included genes in loci BV82_1337, BV82_1618, BV82_2032 (*mexE*), BV82_3631 (*mdtA_1*) and BV82_0511 (*czcA*), all forming a network in STRING analysis (Fig. 4). The exudates of maize, but not tomato, caused a strong upregulation of *mexE* (4.12 log_2_FC). BV82_1337 and BV82_1618 were also upregulated specifically in the maize treatment, while *czcA* and *mdtA* were tomato-induced (Table 1). Efflux pumps of the RND superfamily, together with the so-called Omp proteins, form protein complexes that enable Gram-negative bacteria to export harmful drugs out of the cell. A particular trait of the RND transporters is that, unlike some other efflux transporters, they enable the transport of compounds not into the periplasmatic space but directly into the external medium (87).

#### Cell motility and chemotaxis

Motility in bacteria can be divided into three types: swimming, swarming, and twitching. While swimming and swarming depend on the rotations of the flagella, twitching relies on the extension-retraction movements of the type IV pili (88). In the response of P482 to maize-derived compounds, we observed significant upregulation of BV82_2809 (4.20 log_2_FC) encoding FimV – a protein involved in the assembly of type IV pili (TFP). The type IV pili are structures on the bacterial surface known to be involved in microbial adherence and certain types of movement, such as the social gliding motility in *Myxococcus xanthus* and the twitching motility in *Pseudomonas* and *Neisseria* species. In *Myxococcus*, the action of TFP was found to be under the control of a chemotaxis-like system. It was postulated that the type IV pili are involved in directed cell movement, biofilm formation, guided tissue invasion, and other pathogenesis-related events (reviewed in: (89)).

The response of P482 to the exudates of maize also involved upregulation of *fliS* encoding flagellar protein FliS (BV82_0871; 1.87 log2FC). Such upregulation was not observed in contact with tomato-derived compounds. Moreover, in the case of tomato, we saw upregulation of BV82_3459 (2.22 log_2_FC) encoding a protein presumed to be a regulatory helix-turn-helix LysR family protein that negatively regulates the transcription of the flagellar master operon *flhDC* (90). It was shown that although flagella are essential for some *Pseudomonas* strains to colonize a particular host, their presence may also lead to disadvantages related to the onset of defense responses by the flagellar antigens (91, 92).

A process related to motility is chemotaxis. Tomato exudates caused downregulation of two genes encoding methyl-accepting chemotaxis signaling domain proteins (MCPs) in loci BV82_3217 and BV82_3844. The MCP proteins are the predominant chemoreceptors in bacteria and archaea. However, they are more abundant and versatile in bacteria that inhabit dynamically changing environments and in bacteria that are motile. The MCPs were found to be involved in regulating many microbial activities, including, but not limited to, biofilm formation, flagellum biosynthesis, and degradation of xenobiotic compounds (reviewed in (93)). Most MCPs described so far originate from the rhizosphere of *Pseudomonas* spp. enable the detection of amino acids, polyamines, or organic acids used as energy sources. The few strains which have been analyzed for the number of MCP genes in the genome were shown to harbour between 27 and 37 MCP genes (94).

#### Copper tolerance

When comparing the transcriptomic response of P482 to the exudates of tomato and maize, we saw a significant difference in the expression of *copA* (BV82_2904) and *copZ* genes (BV82_2906). Although some induction of *copA* and *copZ* was observed in response to tomato (1.50 and 1.97 log_2_FC, respectively), the expression of the two genes was significantly higher in response to the exudates of maize (3.69 and 3.94 log_2_FC, respectively). CopA is a transmembrane ATPase responsible for the efflux of Cu^+^ ions delivered to it by a cytoplasmic chaperone CopZ. Both proteins are the elements of a well-characterized Cu^+^ tolerance systems (95). In P482, an expression pattern consistent with the one observed for the *copA* and *copZ* also occurred in the case of a neighbouring gene BV82_2907 (*bcr*), encoding a drug resistance transporter with homology to the Bcr/CflA subfamily of transporters.

### Verification of RNAseq data by real-time RT-qPCR

To assess the reliability of RNAseq data, results obtained with this method were verified for a selected pool of genes using reverse transcription real-time qPCR (RT-qPCR). The selected targets included: *lrp* (BV82_3254)*, mexE, norC, ssuC, trpB and ytfE*. *lrp* (BV82_3254) – a putative transcriptional regulator; *mexE* – an efflux transporter of the RND family; *norC* – nitric oxide reductase subunit C; *ssuC* – transporter protein involved in the import of sulfur-containing compounds; *trpB* – tryptophan synthase, beta subunit (BV82_2326); *yftE* – hemerythrin HHE cation binding domain protein. The objective was to choose a set of genes representing both upregulated and downregulated genes, with both high and low changes in the expression levels observed between treatments.

Overall, the gene expression profile for all genes tested by RT-qPCR corresponded to the one obtained by RNAseq, confirming the reliability of the RNAseq data (Fig. 5). For some genes like *ssuC* and *trpB* the results obtained by the two methods were near-identical. For *lrp*, for which the worst correlation could be claimed, the expression profile showed the same tendency of up- and-down regulation in the studied treatments, yet the absolute mean log_2_FC values differed depending on the method (RNAseq vs RT-qPCR).

**Fig. 5.**
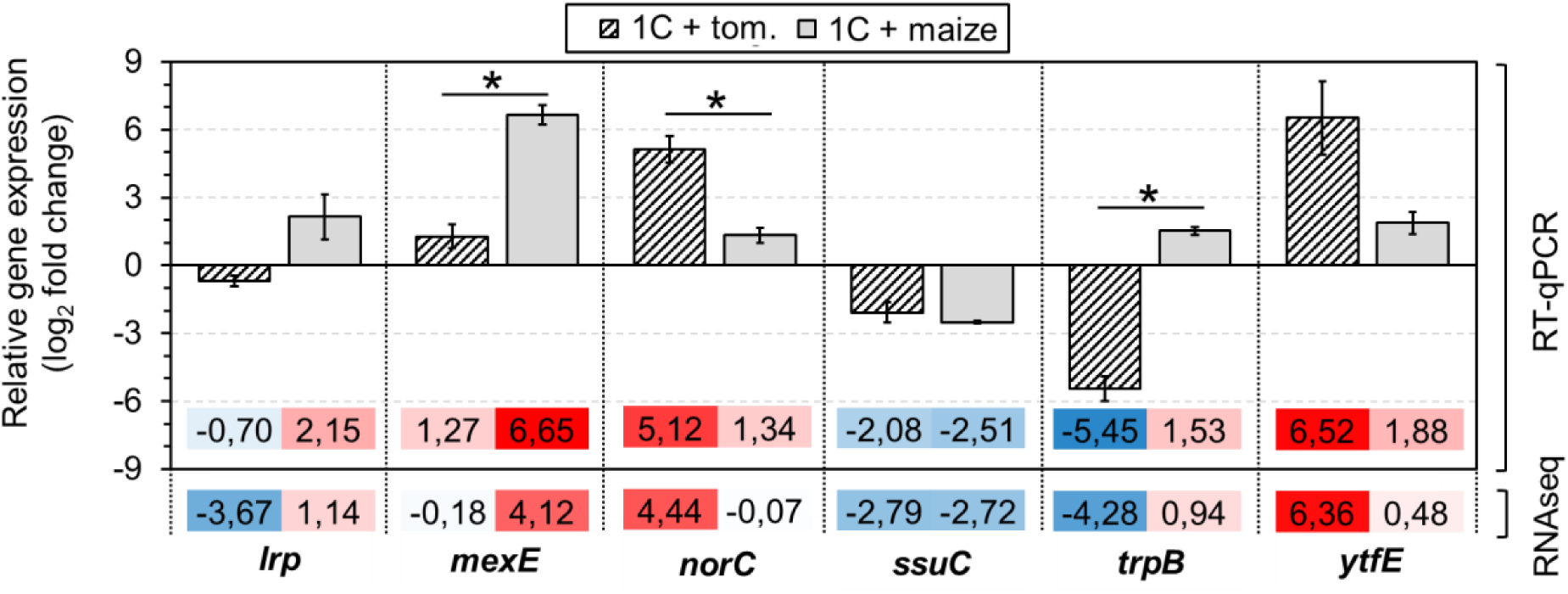
Comparison of gene expression results obtained by RNAseq and RT-qPCR. The RT-qPCR experiment aimed to confirm the reliability of the RNAseq data. Analysis was performed for a group of genes representing both low and high differences in expression between treatments. The investigated targets included: *lrp* – putative transcriptional regulator (BV82_3254); *mexE* – efflux transporter, RND family, MFP subunit (BV82_2032); *norC* – nitric oxide reductase subunit C (BV82_3246); *ssuC* – binding--dependent transport system inner membrane component family protein (BV82_1676); *trpB* – tryptophan synthase, beta subunit (BV82_2326); *yftE* – hemerythrin HHE cation binding domain protein (BV82_3239). ‘1C + tom.’ – P482 grown in 1C with the addition of tomato exudates; ‘1C + maize’ – P482 grown in 1C with the addition of maize exudates. qPCR results were normalized to two reference genes: *gyrB* and *rpoD*. All results were scaled to samples form P482 grown in 1C medium alone. Bars represent mean values for 3 biological replicates. Error bars show standard error of the mean (SEM). Stars indicate statistical significance of differences between groups (p<0.05). Blue and red backgrounds indicate down- and up-regulation of gene expression, respectively.

### Differences in the chemical composition of exudates

Non-targeted analysis of the chemical composition of the samples was performed using GC-MS and ^1^H-NMR. Detailed description of the results is provided in SI Appendix (includes Table S8, Fig. S5, Fig. S6). A difference in the composition of exudates that could be linked to other results is the content of glucose. While glucose was highly abundant in the exudates of maize (9947 ng mg^-1^), this sugar was not detected in the tomato-derived samples.

Additionally to GC-MS and ^1^H-NMR, we applied Liquid Chromatography-Selected Reaction Monitoring (LC-SRM) to assess the relative quantity of 21 amino acid in the exudates of both plants. In particular, we were interested in differences in the content of tryptophan, phenylalanine, methionine, valine, leucine and isoleucine, as genes involved in the metabolism or response to those amino acids were among GDRs. The content of branched chain amino acids valine and leucine in maize exudates was approx. 2 log_2_fold higher (4 times higher) than in tomato, and equal for both samples in the case of isoleucine. This is in contrast with increased catabolism of those amino acids in P482 in the presence of tomato exudates, not maize. The quantity of aromatic amino acids phenylalanine and tryptophan was 2.2 and 1.2 log_2_fold higher in maize than in tomato, respectively. In terms to the related transcriptomic response, in tomato, but not maize, the catabolism of phenylalanine was increased and tryptophane synthesis decreased (Fig. 4). Differences in the methionine levels vere not statistically significant. Results for the remaining amino acids are provided in the SI Appendix.

## Discussion

Two bacterial phyla are universally enriched within the rhizosphere of plants: Bacteroidetes and Proteobacteria (96). In the case of the proteobacterial genus *Pseudomonas*, the ability to thrive in various environmental niches is attributed to the metabolic flexibility of these microorganisms (8). It is yet unclear which particular metabolic changes are required of bacteria promiscuously colonizing plant roots to adapt to different plant hosts, or to stay associated with a given host when the physiological state of thereof is subjected to change. Drawing general conclusions from existing data is challenging as the majority of studies include single strain-host setups. Furthermore, the matter of specificity in plant-microbe interactions is studied in depth for symbiotic rhizobia, but not to a similar extent for phytobacteria that form less intimate associations with their hosts (97).

In this study, we used *P. donghuensis* P482 as a model for a promiscuous root colonizer. The strain was isolated from the rhizosphere of tomato (*Solanum lycopersicum* L.), a Dicot plant, but it can also colonize the rhizosphere of phylogenetically distant maize (*Zea mays*) – a Monocot species (Fig. 1). We performed a comparative analysis of transcriptome changes in P482 upon stimulation with root exudates of tomato and maize. Our goal was to investigate the metabolic shifts undertaken by bacteria during the interaction with different plant hosts, and to identify the host-independent aspects of response to the exudate compounds.

### Exudates of tomato caused more extensive changes in the transcriptome of P482 than the exudates of maize

The response of *P. donghuensis* P482 to the exudates of tomato was more pronounced than to the exudates of maize, with 8.15% and 3.70% of the transcriptome affected, respectively (Fig. 2). Similar results as for P482 and tomato were reported for *P. aeruginosa* PAO1 in response to the exudates of two varieties of beetroot (*Beta vulgaris*), each known to select for different microbial populations (16). Depending on the variety, 9.30% and 8.13% of the transcriptome of *P. aeruginosa* was affected (16). In both studies, our on P482 and of Mark et al. (16) on PAO1, the pool of bacterial genes differentially affected by the tested plant hosts was larger (3-6 times) than the pool of genes of the shared response of bacteria to the exudate compounds.

### Shared response of P482 to root exudates involved downregulation of genes for the acquisition of sulfur, heme, and iron carriers sensed by FecR, as well as the assembly of T6SS and amino acid transport, while genes for arsenic resistance and *bfr* for iron homeostasis were upregulated

Several genes of the *ssu*, *tau* and *cys* operons for sulfur assimilation were downregulated in P482 as part of the host-independent response of P482 to root exudates. Downregulation of the *ssu* genes, or both the *ssu* and *tau* genes, was also observed for 6 out of 8 *Pseudomonas* spp. strains exposed to the root exudates of grass *Brachypodium distachyon* (98). This implies that plant exudates may be an important source of sulfur for the rhizosphere pseudomonads. Interestingly, an opposite situation regarding sulfur supply in plant-microbe interactions was observed in an *Arabidopsis thaliana* – *P. syringae* model during a foliar infection, where plants inflicted sulfur starvation in the pathogen to limit its virulence (99).

The highest upregulation among the genes of the shared response of P482 to exudates was observed for *arsC,* encoding an arsenate reductase, and *arsH,* encoding an organoarsenical oxidase (40, 41). Arsenic is a highly toxic metalloid ubiquitous in the environment. Its two inorganic forms are arsenite (As(III)) and arsenate (As(V)). As(III) is predominates in anaerobic (reducing) environments while As(V) in aerobic (well-oxidized) conditions (39). Bacteria developed different mechanisms to process arsenic and they play a vital role in the biogeochemical cycle of this toxic compound (100). Moreover, plant-associated, arsenic-resistant bacteria were found to increase the tolerance of soil-derived arsenic in plants (39). Microorganisms were also reported to use methylated arsenicals as weapons against their microbial competitors (101). Interestingly, in the context of the latter, the *arsH* gene, upregulated in P482 in the presence of exudates, was first described as a determinant of microbial resistance to trivalent forms of herbicide monosodium methylarsenate (41).

Gene networking analysis revealed that the arsenic resistance and sulfur assimilation pathways in P482 may be linked (Fig. 4). Relation between arsenite (As(III)) stress and sulfur metabolism was studied in the eukaryote *Saccharomyces cerevisiae*, in which sulfate assimilation and glutathione biosynthesis pathways were induced upon arsenite exposure (102). Correlation between sulfur metabolism and arsenic resistance was also reported in the procaryotic organism *Herminiimonas arsenicoxydans,* an ultrasmall bacterium isolated from industrial sludge (103). In *H. arsenicoxydans*, early stress induced by arsenite (15 minutes post-exposure) strongly stimulated the expression of genes related to sulfur assimilation, sulfur oxidation and glutathione synthesis. However, the upregulation of sulfur-related genes was not observed in what the authors termed a ‘late response’ to arsenite stress, that is at 6 and 8 hours post treatment (104). In the strain P482, as part of the shared response to root exudates, we observed strong upregulation of *arsC*, as well as *arsH*, with a simultaneous downregulation of genes of multiple sulfur acquisition pathways. This is in contrast to the abovementioned studies where arsenic resistance and sulfur assimilation pathways had a tendency for simultaneous upregulation. Based on the current study, we have yet no way to state if the two pathways in P482 are truly connected and how.

Iron is a trace element required for the functioning of cells. Its bioavailable fraction in the environment is low, therefore bacteria secret iron-scavenging molecules called siderophores. *Pseudomonas* spp. are known to produce multiple siderophores, with fluorescent pyoverdine being most characteristic for this group (105). *P. donghuensis* strains also produce 7-hydroxytropolone (7-HT) – a compound with iron-scavenging and antimicrobial properties (21, 106, 107). Some microorganisms can use exogenous siderophores, therefore shifting the energetic cost of their production to other organisms (108). This phenomenon is sometimes referred to as ‘sierophore piracy’ (109). In *P. donghuensis* P482, the response to exudates of both tomato and maize involved the downregulation of seven genes related to the acquisition of ferric citrate and/or potentially other iron-binding xenosiderophores, as well as the downregulation of selected *has*, *hem*, and *hmu* genes involved in the acquisition of heme. The expression of genes encoding pyoverdine and 7-HT was not altered when compared to the medium-only control. The obtained results suggest a lowered demand for iron sequestration by P482 in the presence of root-collected compounds. In line with what we observed in P482, the expression of pyoverdine was also not upregulated in any of the eight tested *Pseudomonas* strains in response to the exudates of *B. distachyon* (98). In fact, upregulated expression of gene(s) somehow related to the acquisition of iron was reported only in four out of eight tested pseudomonad strains. Those genes encoded proteins of iron dicitrate transporters (*Pseudomonas* spp. strains SBW25 and 30-84), genes for the biosynthesis of siderophores ornicorrugatin (strain SBW25) or pyochelin (strain Pf-5), heme-degrading enzymes (strains 2– 79, 30–84), and TonB-associated genes (strains 2–79 and Pf-5) (98). It is also interesting to discuss the obtained results with the work of (17). These authors compared the transcriptome of *P. protegens* CHA0, a plant-associated strain with insecticidal properties, during the interaction of the bacterium with wheat and two insect hosts. They found that the expression of genes related to heme acquisition and pyoverdine synthesis in CHA0 was relatively low in response to wheat when compared to the high expression observed during the infection of insects *Plutella xylostella* and *Galleria mellonella*.

Plants were also found to produce siderophore-like compounds. Although the presence of iron-helating mugineic acid was also detected in Dicot plants (110), the phenomenon is best known to occur in Graminaceous plants (i.a. grasses). Boiteau et al., (108) have found that the production of phytosiderophores was increased in *B. distachyon* following inoculation with *P. fluorescens* SBW25 compared to the amount in sterile roots. Moreover, in response to Fe deficiency, the *P. fluorescens* consumed the plant-derived siderophores present in the exudates rather than producing its own (108). It therefore seems that rhizosphere-associated pseudomonas generally do not suffer from iron starvation when supplied with root exudates. However, a greater effort in terms of iron acquisition could be required in the rhizosphere in the presence of microbial competitors (51). Production of siderophores was shown to be an important mechanism of biocontrol, that is the ability of root-associated microbiota to suppress the activity of plant pathogens (111).

Presence of other (micro)organisms could also have an effect on the activation of the type VI secretion system (T6SS), the assembly of which was downregulated in the presence of root exudates in P482 monocultures. T6SS is a nanomachine used by bacteria to inject effectors to other procaryotic or eucaryotic cells (112). It is known for its role in interbacterial competition and was shown to protect PGPR from eukaryotic bacterivores (51).

Contrary to downregulating genes for iron acquisition, root exudates of both tomato and maize significantly upregulated the expression of bacterioferritin gene *bfr* for intracellular iron homeostasis. Bfr protein enables safe storage of iron which, when present in excess in a free form, can cause oxidative stress in the cells through generation of reactive oxygen species (113). Bacterioferritin and Bfr-interacting proteins were shown important for stress resistance and virulence in both plant and animal pathogens (114, 115). They also play a role in the root-nodulating plant symbionts (116). In a recent proteomic study by (117), the root exudates of intercropped maize were reported to upregulate the expression of the bacterioferritin comigratory protein in nodule-forming rhizobacterium *Ensifer fredii*.

### P482 cells treated with tomato exudates switched to nitric oxide detoxification, repair of iron-sulfur clusters, respiration through the cyanide-insensitive cytochrome *bd*, and activation of the catabolism of amino acids and/or fatty acids

Comprehensive analysis of the tomato-specific response of P482 to exudates led us to conclude that the cells are under nitrosative stress. We observed upregulation of genes encoding the Hmp flavohemoglobin, nitric oxide reductase Nor, and the nitric oxide stress protein NnrS (57, 58, 67). Nitric oxide is a freely diffusible molecule that exerts toxic effects on bacterial cells. To counteract nitrosative stress, bacteria have evolved several pathways to convert NO to non-harmful molecules like N_2_O, NO_3_^-^ or ammonia (118). Efficient prevention of the NO-mediated damage is critical in the case of denitrifying bacteria, which generate NO as an intermediate during respiration, and in microorganisms that need to colonize host tissues, the latter requiring coping with NO produced as a part of the host defenses during infection (119). Deleterious effects of NO result predominantly from the inactivation of proteins containing iron-sulfur (Fe-S) clusters (120, 121). Iron sulfur clusters are redox-active protein cofactors present in almost all organisms and required for many fundamental biochemical processes (89). In P482, we observed upregulation of proteins involved in the repair and the assembly of Fe-S clusters alongside those directly involved in NO detoxification. This included very strong upregulation of *yftE*, known to be a part of bacterial defense against nitric oxide (66), as well as *iscR*, *iscS*, *iscU*, *iscA*, *hscB*, *hscA, fdx* genes related to sulfur cluster biogenesis (70). YftE protein was shown to contribute to the survival of *Yersinia pseudotuberculosis* in spleen following nitrosative stress and contribute to the virulence of this human pathogen (121).

*Pseudomonas* bacteria, under the conditions of oxygen shortage, can perform denitrification – a process of using nitrate as an alternative acceptor of electrons (122). Denitrification involves a stepwise reduction of nitrate to N_2_, along which a variety of intermediate compounds are produced, including nitric oxide (NO). In *P. donghuensis* P482 grown in the presence of tomato exudates, the upregulation of NO-detoxifying genes was not accompanied by the induction of *nir* genes, the latter of which would be indicative of an active denitrification process (60). This implies that the strong upregulation of the activity of NOR is not related to increased indigenous production of NO due to a switch of P482 to a denitrification pathway, but rather is aimed at coping with the toxicity of exogenous NO, here derived from the tomato exudates.

Plants use NO as a signal molecule involved in a diverse range of physiological processes including germination, development, flowering, senescence, as well as the response to biotic and abiotic stress. Considering NO is highly reactive, it is stored in the form of S-nitrosothiols (SNOs) which act as reservoirs of NO *in vivo* (reviewed in (123)). NO plays an important role in the response of the plant to microbial pathogens, but it was also shown in the establishment of symbiotic interactions between rhizobia and legumes (65). Interestingly, in an animal model, NO is made by host tissues during colonization of the squid light organ by *Vibrio fischeri*, what is presumed to help in excluding nonsymbiotic bacterial species (124). Noteworthy, the apparent presence of nitrostive stress in P482 in the presence of tomato-derived compounds did not negatively influence the growth rate of the bacterium *in vitro*. Supplementation of 1C medium with increasing concentration of tomato exudates positively correlated with a steeper growth curve and increased end-point cell density (SI Appendix, Fig. S3). This implies that P482 is well-adapted to coping with the encountered conditions.

Upregulation of *aceA* in P482 in the presence of tomato exudates indicates activation of the glyoxylate shunt – a two-step pathway that bypasses the CO_2_-producing steps of the tricarboxylic acid cycle (TCA). Genes of glyoxylate cycle were also activated in *Salmonella* strain grown in the presence of lettuce exudates (125). In many bacteria, including *Pseudomonas* spp., the shunt is activated when acetate or fatty acids are the primary sources of carbon and energy available to the cell, to counter the problem of recycling citrate to oxaloacetate when acetyl-CoA is the only available carbon source (reviewed in: (80) and (81)). Apart from its essential role in acetate and fatty acid metabolism, glyoxylate shunt may play a still poorly explored role in stress defense and pathogenesis, as activation of the pathway was reported under conditions of oxidative and antibiotic stress, as well as during host infection (81). The expression of *aceA* in *P. aeruginosa* was shown to be upregulated under H_2_O_2_-induced oxidative stress and under iron-limiting conditions, and to be tightly linked to intercellular iron homeostasis (126). This metabolic pathway is also centrally important for the growth and virulence of *P. aeruginosa* during infection scenarios, when the bacterium shows a particular predilection for the catabolism of fatty acids (127). Moreover, shunting of the TCA, together with the shift toward the utilization of amino acids, was shown to increase antibiotic resistance in *P. aeruginosa* from patients suffering from cystic fibrosis (128).

Activation of glyoxylate shunt in P482 in the presence of tomato-derived compounds was accompanied by an increase in the conversion of alpha-keto acids to CO_2_, the induction of the overlapping pathways of branched chain amino acids (BCAA) and fatty acid catabolism, as well as the catabolism of phenylalanine, glycine and, possibly, methionine. In *P. aeruginosa*, the catabolism of BCAA was shown to contribute to the pathogenesis of the bacterium in *Caenorhabditis elegans* through manipulation of host energy status and mitochondrial stress signaling potential (129). As oppose to the situation described for P482 and *P. aeruginosa*, in *S. typhimurium* and *E. fredii* exposed to the root exudates of lettuce and intercropped maize, respectively, upregulation of BCAA synthesis rather that their catabolism has been reported (117, 125). For some pathogenic bacteria it was shown that a balance between complying to the BCAA nutritional requirement and BCAA starvation supports not only proliferation during infection, but also the evasion of host defenses (130). Interestingly, relative analysis of amino acid content of exudates in this study showed that BCAA were more abundant in the exudates of maize than in the exudates of tomato, despite their catabolism was upregulated in the latter (SI Appendix, Table S11).

When it comes to metabolism of amino acids or fatty acids, the role of the involved proteins may be broader than the initial assumptions. For example it was postulated that *P. fluorescens* Pf-5 uses existing pathways for the catabolism of fatty acids, including *mmgC*, during the degradation of toxic compounds like naphthenic acids (131). Other studies showed an additional function the of the *phhA* gene, encoding the bacterial phenylalanine 4-hydroxylase (PAH). In *P. fluorescens,* PAH was shown to be involved not only in the catabolism of phenylalanine, but also in the biosynthesis of melatonin from tryptophan (132, 133). Melatonin acts as an antioxidant which promotes resistance to oxidative stress but it can also function as a growth promoting regulator in plants (reviewed in (134)). Induction of homologues of both *mmgC* and *phhA* was observed in P482 in the presence of tomato exudates.

Tomato-derived compounds caused a significant downregulation of tryptophan synthesis in P482. Contrary, the synthesis of methionine was increased, as could be inferred from the significantly upregulated expression of *metE*. Genes related to methionine synthesis were found to be upregulated in *E. coli* following treatment with S-nitrosoglutathione in conditions mimicking nitrosative stress. In the latter study, *E. coli* mutants with inactivated genes of the methionine synthesis pathway showed decreased tolerance to S-nitrosoglutathione and this effect could be abrogated by the addition of exogenous methionine (59). Intriguingly, the requirement for methionine synthesis was found essential for establishing symbiosis between nitrogen-fixing *Sinorhizobium meliloti* and its plant host *Medicago sativa* (alfalfa) (135). Interestingly, methionine synthesis was decreased in PAO1 in response to exudates of beetroot (16) and in *S. typhimurium* in response to the exudates of lettuce (125).

In P482, in the presence of tomato exudates, we saw downregulation of genes encoding the *cbb3*-type cytochrome c oxidase, while genes *cioA* and *cioB* encoding cytochrome *bd* (CIO) were significantly upregulated. Pseudomonads harbor a branched respiratory chain. Five different terminal oxidases are known to operate in *P. aeruginosa*, allowing the bacterium to exploit an electron transport chain best suited under given circumstances (61). Terminal oxidases *bo_3_* (Cyo) and the cyanide-insensitive oxidase CIO receive electrons directly from ubiquinone. Transport of electrons from ubiquinone to the three remaining terminal oxidases: *aa3*, *cbb_3_* oxidase 1 and *cbb_3_* oxidase 2 is mediated via the cytochrome *bc_1_* complex and c-type cytochromes or a small blue-copper protein azurin (61, 126). The *cbb_3_*-type oxidases, downregulated in P482 in the presence of tomato exudates, are heme-copper oxidases with high affinity to oxygen that were found to play a particular role in cell survival under microaerobic conditions (136). Meanwhile, the upregulated *cio* genes were originally associated with bacterial resistance to hydrogen cyanide, yet there are also other factors that can affect their expression. Cyanide-insensitive cytochrome *bd* was also reported to have a role during copper limitation under aerobic conditions (71) and in conferring tolerance to both oxidative and nitrosative stress (137). For example, it was found to augment the defenses of *Salmonella enterica* against NO in systemic tissues (138). Moreover, in *Pseudomonas* sp. WCS365, *cioA* was found to decrease the fitness of the bacterium in an immunocompromised mutant of *A. thaliana* but not in a wild-type plant, implying the role of this gene in evading host defenses (139). It can be speculated that the reason for the rerouting of the respiratory chain in P482 towards cytochrome *bd* (CIO) is the inhibition of cytochrome complex *bc_1_*, required to pass the electrons to the cytochrome c oxidases including *cbb_3_*, but not to CIO. Such compensatory mechanism was observed in *Mycobacterium tuberculosis* in the presence of inhibitors of the *bc_1_* complex – an effect which was greatly enhanced by simultaneous addition of NO donors (140). It is also important that cytochrome *bd*, though being inhibited by nitric oxide, exhibits an unusually fast NO dissociation rate from its active site (141).

### Maize-specific response to exudates included upregulation of genes associated with motility, the expression of MexE and two other RND efflux pumps, and copper tolerance

Maize-specific response of P482 to exudates involved upregulation of the FimV gene related to the assembly of type IV pili and *fliS* encoding flagellar protein FliS. Flagella are known for their role in in bacterial swimming motility, while type IV pilli are involved in directed cell movement by twitching or gliding, as well as in biofilm formation, guided tissue invasion, and other pathogenesis-related events (reviewed in: (142)). Type IV pilli were found essential for endophytic rice colonization by the N2-fixing endophyte *Azoarcus* sp. strain BH72 (143). Interestingly, gene *pilK*, involved in twitching motility, was downregulated in *P. aeruginosa* in response to the exudates of beetroot (16). The tendency for expression of motility-related functions in P482 in the presence of tomato exudates was the opposite than in the response to maize, that is towards their suppression. This could be inferred from the upregulation of a negative regulator of the flagellar master operon *flhDC* (90) and the downregulation of two genes encoding methyl-accepting chemotaxis signaling domain proteins (MCPs), presumably involved in chemotaxis (reviewed in (93)).

Flagella were shown essential for some *Pseudomonas* strains to efficiently colonize their host. *P. fluorescens* F113 with a mutated *fliS* gene was found to be non-motile and unable to compete with the wt strain for the colonization of the root tip of alfalfa (91). Motility was also shown to increase the infectivity of two pathogens, the *P. phaseolicola* (currently *P. syringae* pv. *phaseolicola*) and *P. syringae* pv*. glycinea*, in the leaves of bean and soybean, respectively (144, 145). On the other hand, a non-motile mutant of *P. syringae* pv*. tabaci* was able to propagate and cause disease-like symptoms on tomato (not an original host), indicating that motility is not required for the survival of this pathogen on this particular plant (146). It was also shown that although flagella can provide advantages to the cells, their synthesis may also lead to some disadvantages. Recognition of certain flagellar antigens may induce hypersensitive response and cell death as part of the host’s immune response (reviewed in: (146)). A decrease in flagella synthesis is considered a major mechanism in the evasion of plant immunity (147). Moreover, studies on *P. putida* KT2440 concerning the metabolic tradeoffs of producing flagella showed that a non-flagellated mutant strain presents a shorter lag phase and is more resistant to oxidative stresses than the *wt* strain. It was suggested that lack of the metabolic expense on producing flagella might provide cells with a surplus of energy (ATP) and reduce power (NADPH) that the cells can allocate for other activities, including stress resistance (92). It remains to be elucidated if the differences in expression of motility genes in P482, observed between tomato and maize treatments, are related to the varying importance of those genes in the colonization of these two hosts, or maybe rather the stress conditions imposed by tomato exudates made the expression of those genes unfavorable.

The exudates of maize caused upregulation of three genes linked to RND-type efflux pumps. In particular, there was a strong upregulation of *mexE*. Efflux pumps of the RND superfamily, together with the Omp proteins, form protein complexes that enable Gram-negative bacteria to export harmful drugs directly to the external environment, and not, like in the case of other efflux systems, into the periplasmatic space. This makes RND efflux pumps important determinants of multidrug resistance, both intrinsic and elevated (87). In *P. aeruginosa,* the intrinsic resistance to antibiotics is related to the presence of several efflux operons: *mexAB-oprM, mexAB-oprM mexCD-oprJ* and *mexEF-oprN*. The latter operon, containing the *mexE* gene strongly upregulated in *P. donghuensis* P482 in the presence of maize exudates, in *P. aeruginosa* is positively regulated by a protein belonging to the LysR family of transcriptional activators (*mexT*) and confers resistance to quinolones, chloramphenicol, and trimethoprim (148). Quinolone resistance based not on modification of the drug target but the activity of an efflux pump was studied in a root-colonizing strain of *Stenotrophomonas maltophilia* (149). In this strain, the respective pump is encoded by the *smeDEF* operon under the control of SmeT repressor. Interestingly, based on the annotation of genes in NCBI database, the SmeDEF pump belongs to the RND family of efflux transporters, just as the *mexE*-containing operon in P482. The authors investigating the role of SmeDEF in *S. maltophilia* considered that it is unlikely that the primary function of this efflux pump, evolved over millions of years, would be resistance to quinolones-synthetic antibiotics present in the natural environment only since the last few decades. Their study shows that the expression of *smeDEF* is triggered by plant-produced flavonoids and that inactivation of the pump by deletion of *smeE* impairs the ability of *S. maltophilia* to colonize the roots of the oilseed rape plant (149).

Upregulation of *mexE* and two other genes related to RND transporters was specific for the response of P482 to the exudates of maize. However, there were also two other genes linked to this family of efflux transporters which were upregulated in response to tomato. What is interesting about this family of efflux pumps is that, in *P. donghuensis* P482, inactivation of gene BV82_4243 encoding another RND-type transporter resulted in a reduced antibacterial activity of the mutant strain against *Dickeya solani*, as well as in the lack of export of the fluorescent siderophore pyoverdine (44). This implies that proteins from the family of RND-type transporters may be involved not only in defence against potentially harmful chemicals, presumably involving secondary metabolities of plants, but also in the export of the strain’s own metabolites.

Last but not least, we observed significantly higher upregulation of *copA* and *copZ* in the response of P482 to maize exudates compared to tomato, qualifying the genes into the differential response pool. Copper is an essential trace element for all organisms, however it is toxic in excess (150). Mavrodi et al., (98) showed considerable upregulation (approx. 5-7 log_2_FC) of one or more *cop* genes related to copper tolerance in 5 out of 8 investigated pseudomonads treated with the exudates of *B. distachyon* (98). Both maize and *B. distachyon* are Monocot grasses.

### Transcriptomic response to exudates through the prism of simplified experimental setups

The coevolution of plants and microorganisms led to the formation of complex networks of interactions between the two groups (5, 151). Mimicking full spectrum of these interactions in the laboratory remains unattainable. What is more, it is technically challenging to study bacterial transcriptome *in planta*, due to the difficulty in isolation of bacterial RNA from complex plant-based matrixes where bacterial cells are present in small numbers (152). In the light of those challenges, the influence of exudates on P482 was assessed for bacterial cells were grown *in vitro* in a minimal medium supplemented with water-collected plant exudates and glucose. A similar approach was applied by (98) in the study on the impact of exudates of *B. distachyon* on different *Pseudomonas* spp. Other media used in studies concerning the influence of exudates on bacteria include casamino acids medium (CAA), applied to investigate the impact of beetroot exudates on PAO1 (16), and Lysogeny Broth (LB), applied to study the response of *Bacillus mycoides* to the exudates of potato (153). In the discussed types of setups, bacterial response to plant-derived compounds may be distorted by the presence of the additional carbon/nitrogen source(s). In the case of P482, chemical analysis of exudate composition revealed that glucose, used in our setup as the single added carbon source for increasing cell yield, was originally present only in the exudates of maize. The time point of sampling may also be of importance. The cells of P482 were harvested upon transition to stationary phase, therefore at a timepoint where the less abundant carbon sources originating from exudates, used with preference, may have already been exhausted. The latter may be a reason why we did not detect clear-cut connections between the differential response of P482 to the exudates of tomato and maize and the differences in the chemical composition of the exudates of these species, at least in terms of simple compounds that could act as carbon sources.

### Conclusions and outlook

Understanding plant-microbe interactions is a key requirement for harnessing their potential to support plant health. Methods based on Next Generation Sequencing, including RNAseq, are powerful tools to gather the necessary insight. In an RNAseq-based study for *P. aeruginosa* published in 2005, the majority (≈70%) of bacterial genes affected by the exudates of beetroot encoded proteins with only putative functions or hypothetical proteins (16). Here, in the case of P482, the same was true for only ≈24% of genes with expression altered by the exudates, showing the progress in the development of databases and annotating algorithms. Tools such as the KEGG database for metabolic pathways (154) and STRING for predicting protein-protein interaction networks (30) provide support that cannot be overestimated in the analysis of omics data, including RNAseq. However, it remains challenging to capture the complete picture of metabolic activity of a given microorganism from transcriptomic data, in particular in the case of non-model microorganisms. This may lead to superficial data analysis and new mechanisms underlying plant-microbe interactions may be overlooked. We hope that in the future, through further development of machine learning, it will be possible to overcome the current bottleneck of converting omics data into biologically meaningful mechanistic insights (155). It may yet require insight from the information technology (IT) specialists if our current way of designing experiments and reporting their results is best suited to aid this transition.

In this study, we identified several metabolic pathways differentiating between the response of *P. donghuensis* P482 to the exudates of tomato and maize. We cannot be certain which differences can be attributed to the particular plant species and which are more connected to the physiological state of the plants. The composition of root exudates is known to be influenced by both factors (3). However, we can say that the identified metabolic pathways are involved in the adaptation of *P. donghuensis* P482 to the conditions it can encounter when interacting with these two hosts. Some of the pathways we identified in P482 were already mentioned in terms of interactions between plants and microorganisms in other studies. The fact that some of them, including the catabolism of branched chain amino acids and methionine synthesis, appear to be affected in different manner depending on the investigated model suggests they may be good targets for future research on host adaptation in promiscuous root colonizers. It is also interesting to what extent the ability of plant-associated bacteria to resist plant-inflicted stress is the foundation of successful plant-microbe relationships.

## Author Contributions

DMK coordinated the experimental work, participated in the collection of exudates and plant cultures, processed the exudates for analyses, carried out bacterial cultures, RNA isolation, RT-qPCR, root colonization experiment, functional enrichment, and networking analyses of the RNAseq data, prepared all figures and tables, and drafted the manuscript. MJ grew plants in gnotobiotic conditions and helped with the collection of exudates. ZK performed NMR measurements. MCP performed GC-MS. KM performed the analysis of amino acid content by LC-SRM. SJ conceived the study, acquired funding, supervised the project, and revised the manuscript.

## Acknowledgments

This research was funded by the National Science Centre (Poland), project No. 2017/25/B/NZ9/00513 to S. Jafra.

