## Supplementary Information_Appendix for "Shared and host-specific transcriptomic response of *Pseudomonas donghuensis* P482 to the exudates of tomato (Dicot) and maize (Monocot) shed light on the host-adaptive traits in the promiscuous root colonizing bacteria"

**This PDF file includes:**

Supplementary text (Materials and Methods, Results)
Figures S1 to S6
Tables S1 to S11
SI References

**Other supplementary materials for this manuscript include the following:**

Dataset S1 Raw data from DE analysis (xlsx)
Dataset S2 BioVenn analysis gene lists (xlsx)
Dataset S3 Comparison of gene expression for tomato and maize in SRGs, GDRs, tiDEGS, miDEGs (xlsx)
Dataset S4 DEGs filtered thresholds and BioVenn groups (xlsx)
Dataset S5 Enriched COGs and KEGGs gene lists (xlsx)
Dataset S6 STRING output enrichment and networking (xlsx)

### SI Materials and Methods

### GC-MS

Samples for GC-MS were prepared as described in (Fan et al. 1997). Twenty mL aliquots of water-collected exudates of tomato and maize were acidified to pH 2 with 0.5 M HCl, freeze-dried and silylated in 60 µL of 1:1 (v:v) acetonitrile:N-tert-Butyldimethylsilyl-N-methyltrifluoroacetamide (MTBSTFA, Acros Organics). Derivatization was performed by sonication in glass chromatographic bottles at 60 °C for 3 h in an ultrasonic bath, followed by overnight incubation at room temperature. GC-MS analysis was performed with GCMS-QP2010 SE (Shimadzu). One µL of each sample was separated on a capillary column DB-5MS (30 m × 0.25 mm i.d.) coated with 0.25 µm film (5%-phenyl)-methylpolysiloxane, bonded and cross-linked, with helium gas as a carrier. The temperature of the column was set to 60 °C for 2 min, followed by 20 °C $min^{-1}$ increments to 150 °C, and then 6 °C $min^{-1}$ to 290 °C. The injection (splitless) temperature, transfer line and ion source were set to 260 °C, 290 °C and 200 °C, respectively. The MS was operated in full scan with a range from 45 to 650 m/z and EI ionization (70 eV). Data analysis was performed in Lab Solutions with NIST 11 Mass Spectral Library.

Relative quantitative analysis was performed based on a four-point standard curve for lactic acid (0.00001-0.01 mg $mL^{-1}$, $R^2$=0.999) (Acros Organics) and, to establish the compound-specific response of the detector, two mixed compound standards, each comprising lactic acid and either nine or five other compounds (SI Appendix, Table S9). All standards were processed and analyzed by GC-MS as described for the samples.

#### NMR

For each of the two plant species, two hundred mL of water-collected exudates were freeze-dried. The lyophilized compounds were suspended in 20 mL of sterile high-purity water and treated with 1 g of sterile Chelex 100 resin (Bio-Rad), in a batch extraction to remove NMR-interfering polyvalent metal ions. The resin was removed by centrifugation (5 min, 8500 RCF), and the pH of the samples was adjusted to ~pH 7 with 0.5 M HCl. The aliquots were freeze-dried again and the dry weight of the plant-derived compounds was established. Each sample was re-suspended in 0.65 mL of $D_2O$ and $^1H$ NMR spectra were obtained with Bruker AVANCE III 500 MHz (Bruker).

#### Liquid Chromatography-Selected Reaction Monitoring (LC-SRM)

_Sample preparation_

The stock solutions of 21 standard L-amino acids (1mg/mL): alanine (Ala, Acros), aspartic acid (Asp, MP Biomedicals), cysteine (Cys, BioShop), leucine (Leu, Acros), tyrosine (Tyr, Acros), phenylalanine (Phe, Acros), ornithine (Orn, Alfa Aesar), arginine (Arg, BioShop), glutamic acid (Glu, MP Biomedicals), glutamine (Gln, Roth), glycine (Gly, Sigma Aldrich), histidine (His, BioShop), lysine (Lys, Acros), methionine (Met, MP Biomedicals), proline (Pro, BioShop), tryptophan (Trp, BioShop), taurine (Tau, Alfa Aesar), serine (Ser, Acros), isoleucine (Ile, SAFC), threonine (Thr, Sigma Aldrich), valine (Val, BioShop) and deuterated leucine (LeuD3, L-Leucine-5,5,5-d3, 99% atom D, Sigma-Aldrich), all of at least ≥ 98.5 purity, were prepared in 0.1 M HCl (Sigma Aldrich) and further diluted with water (LC-MS grade, Merck) before the LC-MS/MS experiments. The aqueous root exudates of maize and tomato samples (100 µL each) were spiked with deuterated leucine standard (Leu D3, Sigma Aldrich, 50 ng/mL in the final sample) and evaporated to dryness using a vacuum concentrator. Then, they were dissolved in 10:90 of mobile phase A (10 mM ammonium formate in water) and B (10mM ammonium formate in acetonitrile) (all LC-MS grade, purchased at Sigma-Merck) and transferred to the LC vials.

*Measurements*

Relative quantification of amino acids in maize and tomato root exudates was performed on QTRAP 6500 triple quadrupole tandem mass spectrometer (MS/MS) (SCIEX) coupled in line with Eksigent LC200 (Eksigent) microLC system, which were controlled by Analyst 1.6.2 software (SCIEX). Five µL of each of the samples were injected in five technical replicates each by the CTC PAL autosampler onto the Reprospher HILIC-A 3 µm 150 x 0.5 mm column (Dr Maisch), where their components were separated using mobile phases A and B and the following gradient program: 0-0.5 min – 1% A /99 % B, 0.5-12min – 1% A to 30% A, 12-14min – 30% A / 70 % B, 14-16 min – 30% A to 1% A, 16-20 min 1% A /99 % B (with column equilibration at 1% A /99% B for 20 min), at the flow rate of 20 µL/min. The eluates from the column were subsequently ionized in the positive ion mode in the TurboV ESI ion source (SCIEX), which was working at 5500 V ion spray voltage, 30 curtain gas, 30 ion source gas 1 and 2, medium collision gas and 300°C temperature. Finally, ionized samples were subjected to the MS/MS analyses in the multiple reaction monitoring (MRM) mode at de-clustering potential (DP), collision energy (CE) and cell exit potential (CXP) individually optimized for each analyte (SI Appendix, Table S10); entrance potential of 10 V, 20 ms dwell time, unit resolution and 5 ms pause between mass ranges. Standard solutions of amino acids and syringe infusion to the mass spectrometer were used for the manual development of two MRM transitions for each amino acid (one qualifier, one quantifier) for the LC-MS/MS method and to determine their compounds parameters, namely, DP, CE and CXP (SI Appendix, Table S10).

*Data analysis*

The .wiff files from LC-MS/MS analysis were exported to the MultiQuant 2.1 software (SCIEX). The following parameters of the MQ4 algorithm were applied for extraction of areas under peaks of each MRM transition: Gaussian smooth 1.0 point, expected RT defined for each amino acid separately, RT half window 30 sec, report the largest peak, minimum peak width 2 point, minimum peak height 0; integration parameters: noise percentage 40 %, baseline subtraction window 2 min, peak splitting factor 2 points. The extracted areas under peaks of the quantifier MRM transitions for each amino acid (SI Appendix, Table S10) were then exported to Excel, where the mean area under peak of 5 technical replicates for each sample was calculated. The log2 fold change in each of the analyzed amino acids in maize versus tomato samples and its p-value, determined in a two-sided unequal variance t-Test, were also calculated using Excel features.

### SI Results

#### Differences in the composition of root exudates of tomato and maize

The composition of exudates of 18 days old tomato and maize plants was analyzed by GC-MS and $^{1}$H-NMR. The obtained chemical profiles were compared to identify significant differences between the studied cultivars of the two plant species.

GC-MS analysis revealed 72 picks in total for tomato in comparison to 63 picks for maize (Fig. S5). Out of compounds successfully identified and quantified, the most abundant were found to be, in the exudates of tomato: malic acid (88 ng $mg^{-1}$ of exudate dry weight), azelaic acid (72 ng $mg^{-1}$), succinic acid (53 ng $mg^{-1}$) and lactic acid (49 ng $mg^{-1}$) (Table 1) and, in the exudates of maize: malic acid (1481 ng $mg^{-1}$), citric acid (175 ng $mg^{-1}$), trans-aconitic acid (135 ng $mg^{-1}$), lactic acid (93 ng $mg^{-1}$) leucine (41 ng $mg^{-1}$) and succinic acid (37 ng $mg^{-1}$).

A limitation of GC-MS is that, within the sample, only compounds which undergo silylation can be analyzed. On the contrary, $^1$H-NMR enables unbiased measurement of whole samples, however yielding complex spectra that can pose interpretational challenges. In this study, $^1$H-NMR analysis of exudate samples confirmed the presence of malic acid and citric acid in both samples, as determined by GC-MS. Moreover, $^1$H-NMR revealed the presence of high amounts of glucose (9947 ng $mg^{-1}$) in the exudates of maize – a sugar not detected in the tomato-derived samples (Fig. S6) (SI Appendix, Table S8). A more general conclusion brought by $^1$H-NMR analysis was that while tomato exudates contained relatively small amounts of multiple compounds, the composition of exudates of maize was less versatile, but the compounds were present in higher concentrations.

Compounds present solely in one type of exudates or in significantly higher amounts were considered among the characteristics of a given plant species. In the case of tomato, this included pimelic acid and azelaic acid. Each mg of dry weight of exudates of tomato contained 16 times more pimelic acid and 10 times more azelaic acid than was detected in the exudates of maize. Compounds more typical for maize were glucose, aconitic acid (not seen in tomato), citric acid, leucine, and phenylalanine. Exudates of maize also contained more malic acid than the exudates of tomato. However, as determined by GC-MS, malic acid was the most highly abundant compound in the exudates of both species (SI Appendix, Table S8).

Relative quantification of 21 amino acids in the exudates of maize and tomato was performed using HILIC LC-MS/MS in MRM mode (here referred to as LC-SRM). The method was developed and optimized for this purpose. Differences in the quantity of amino acid between the analyzed samples were expressed as $\log_2$ fold changes (SI Appendix, Table S11). Fourteen amino acids were more abundant in the exudates of maize compared to the exudates of tomato, out of which differences for 10 were statistically significant ($p<0.05$): alanine, arginine, leucine, lysine, ornithine, phenylalanine, proline, threonine, tryptophan and valine. In turn, 5 amino acids were less abundant in maize exudates than in tomato exudates, of which the difference was statistically significant solely for taurine (SI Appendix, Table S11).

**SI Figures**

**Fig. S1 The growth stage of tomato (A) and maize (B) on day 18 when the exudates were harvested.**

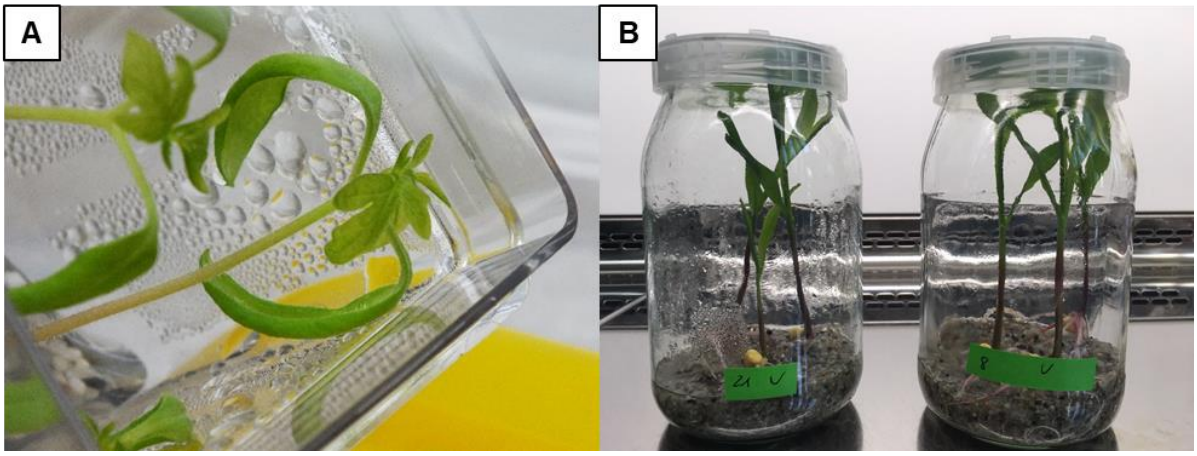

**Fig. S2 PCR products for the corresponding genes following electrophoresis in 1.7% agarose gel.**

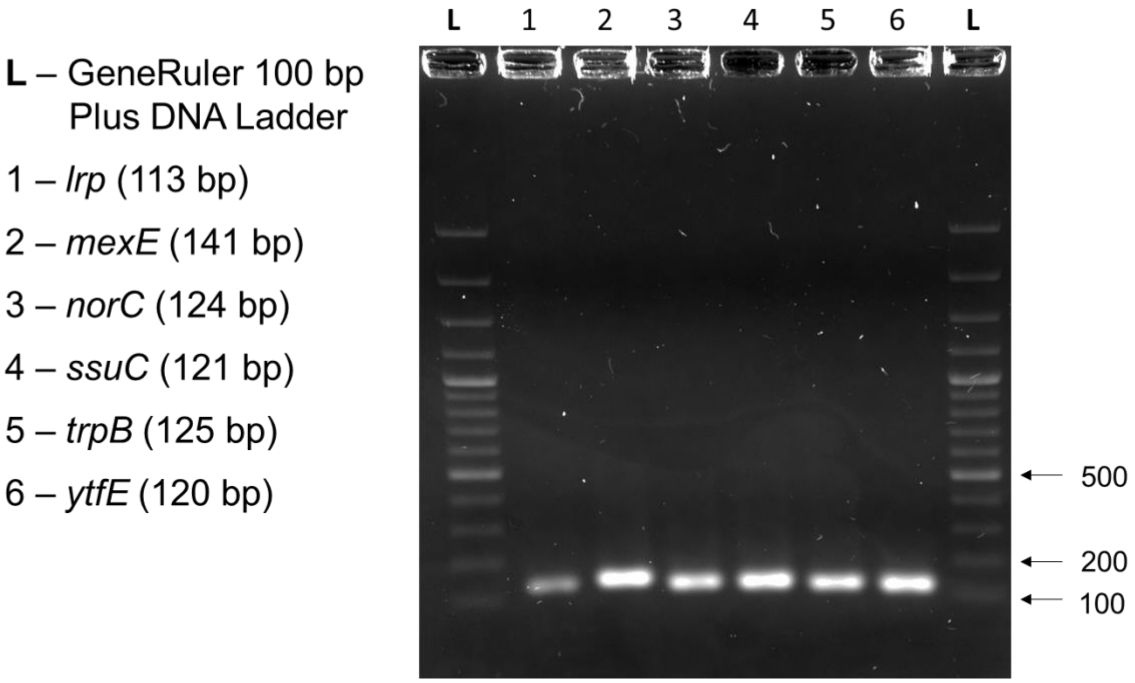

**Fig. S3 Growth of P482 in 1C medium supplemented with root exudates of maize or tomato.** Panels A and B show growth in the presence of different concentrations of the root exudates of maize and tomato, respectively. For the RNAseq experiment, the cells were cultured in 0.2 mg $L^{-1}$. The $OD_{600}$ of cultures was monitored in real-time, and the cells were collected after reaching the early stationary phase per each condition (C).

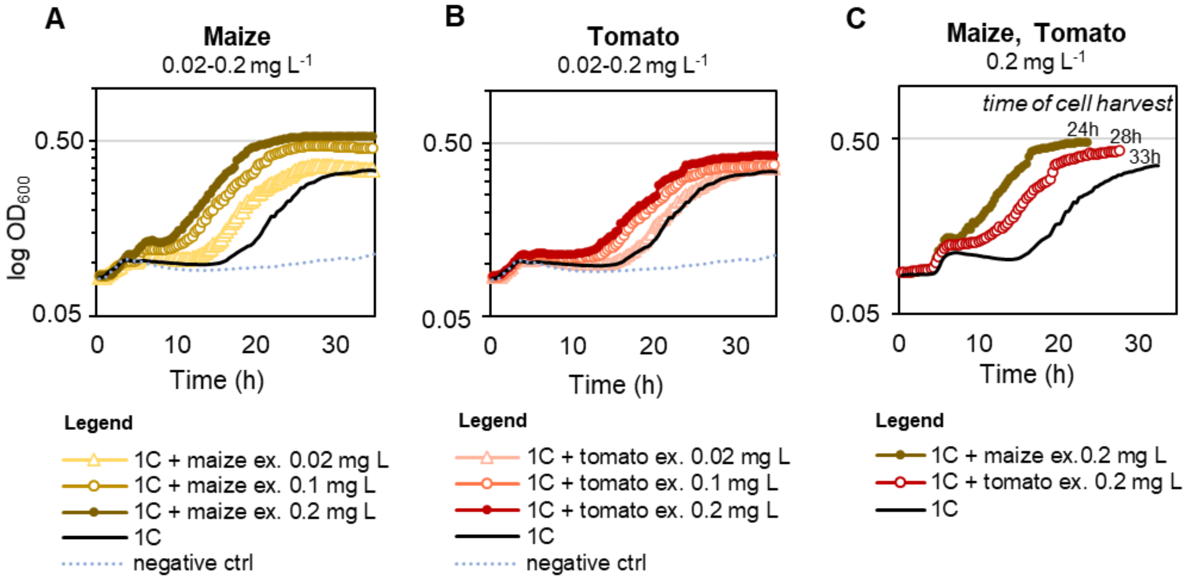

**Fig. S4 Principal component analysis (PCA) plots depict overall transcriptome differences between samples.** PCA assesses the variance in a dataset in terms of components defined on axes x and y. Data points correspond to RNA samples obtained for P482 grown in different experimental conditions: ‘1C’ – unsupplemented 1C medium, ‘1C + tomato ex. 0.2 mg L$^{-1}$’ – medium supplemented with tomato exudates and ‘1C + maize ex. 0.2 mg L$^{-1}$’ medium supplemented with maize exudates. Panels A-C show: A – 1C vs 1C + tomato ex. 0.2 mg L$^{-1}$; B – 1C vs 1C + maize ex. 0.2 mg L$^{-1}$, C – 1C + tomato ex. 0.2 mg L$^{-1}$ vs 1C + maize ex. 0.2 mg L$^{-1}$.

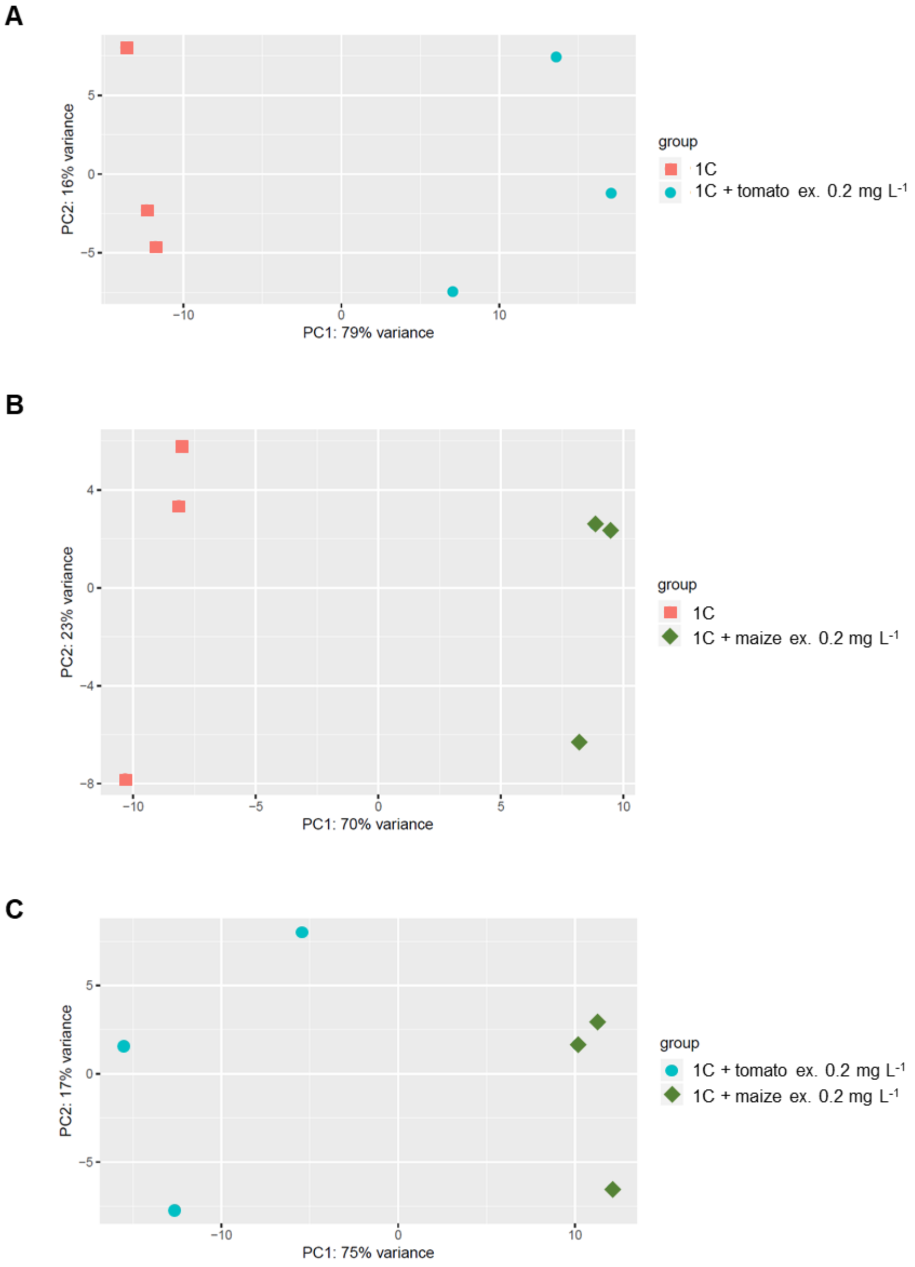

**Fig. S5 Total ion chromatograms obtained by GC-MS for sylilated root exudates of maize cv. Bajm (A) and tomato cv. Saint Pierre (B).** The compounds were identified based on mass spectra.

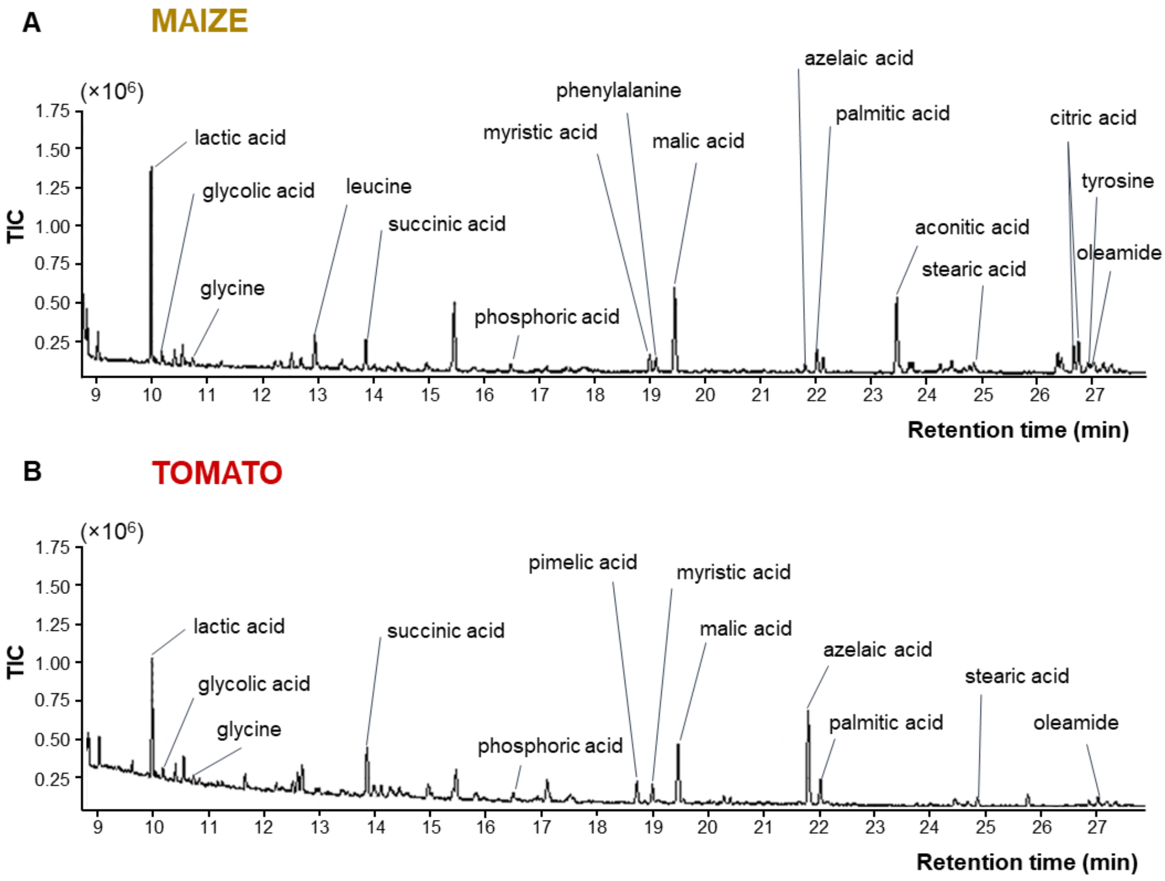

**Fig. S6 1-D $^1$H NMR spectra of maize (A) and tomato (B) root exudates.**

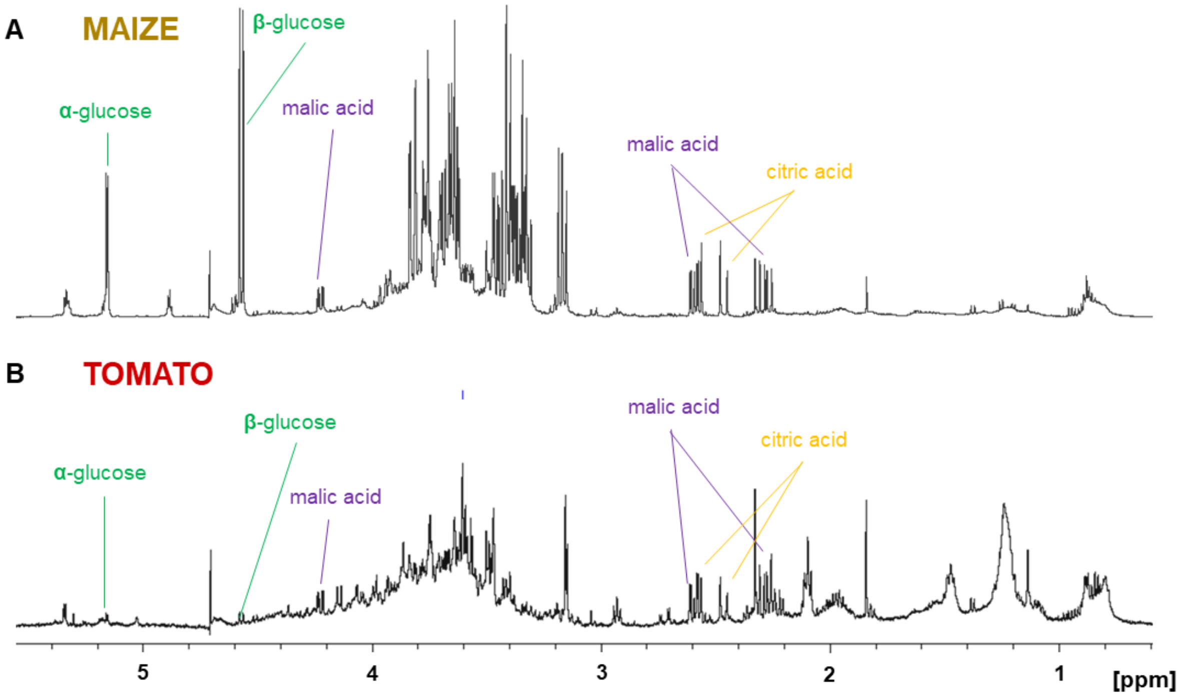

### SI Tables

**Table S1** List of PCR primers used in this study with the established amplification efficiencies.

| **Target gene (locus)** | **Primers** | **Primer sequences** | **Amplicon length (bp)** | **E**[A] | **SE (E)**[B] | **$R^2$** [C] | **Slope** | **Reference** |
|---|---|---|---|---|---|---|---|---|
| *gyrB* (BV82_2296) | F_ gyrB<br>R_ gyrB | 5’ ATCGACAAGCTGCGCTATCA<br>5’ CGGCTGAGCGATGTAGATGT | 144 | 1.96 | 0.08 | 0.981 | -3.41 | (Matuszewska et al. 2021) |
| *rpoD* (BV82_1895) | F_ rpoD<br>R_ rpoD | 5’ CCACGACGGTATTCGAACTT<br>5’ CGTGCCAAGAAAGAAATGGT | 152 | 1.96 | 0.02 | 0.999 | -3.42 | (Matuszewska et al. 2021) |
| *lrp* (BV82_3254) | F_lrp<br>R_lrp | 5’ CCCGAGGTCAACCACAACTA<br>5’ GTGTCGGCTTCCAGTTCGT | 113 | 1.944 | 0.008 | 1 | -3.463 | This study |
| *mexE* (BV82_2032) | F_mexE<br>R_mexE | 5’ GCGTTACCCCCTTGCTCTAT<br>5’ GTGAACTCGTCCCATTCGTT | 141 | 1.956 | 0.012 | 0.999 | -3.432 | This study |
| *norC* (BV82_3246) | F_norC<br>R_norC | 5’ ATGCATGCCTGGATGAAGAT<br>5’ TGATGTTCGAGCTCCATTTG | 124 | 1.999 | 0.013 | 0.999 | -3.325 | This study |
| *ssuC* (BV82_1676) | F_ssuC<br>R_ssuC | 5’ GCACCTTGTTCCCGATTTAC<br>5’ AGAATCACCTGGCGAAACAG | 121 | 1.97 | 0.017 | 0.998 | -3.396 | This study |
| *trpB* (BV82_2326) | F_trpB<br>R_trpB | 5’ AATCGATCATCGGCAAAGAG<br>5’ ATCGAGGAAGTCGTGGAACA | 125 | 2.114 | 0.027 | 0.997 | -3.024 | This study |
| *ytfE* (BV82_3239) | F_ytfE<br>R_ytfE | 5’ GACATGCAGCAGGAACTTGA<br>5’ ATGCTCCAGACGCATAACCT | 120 | 1.976 | 0.015 | 0.999 | -3.381 | This study |

[A] primer pair efficiency
[B] efficiency standard error
[C] coefficient of determination for linear regression calculated for a 7-point, 10-fold dilution standard curve

**Table S2 Titer of P482 Rif in the rhizosphere of soil-grown tomato and maize plants.**

| Experiment | Plant no. | Cell titer Maize [A] | Cell titer Tomato |
|---|---|---|---|
| 1 | 1 | 5.17E+05 | 1.58E+07 |
| 1 | 2 | 1.27E+07 | 1.40E+07 |
| 1 | 3 | 1.50E+04 | 7.83E+06 |
| 1 | 4 | 9.33E+04 | 1.07E+07 |
| 1 | 5 | 1.60E+06 | 1.75E+06 |
| 1 | 6 | 1.33E+04 | 1.90E+07 |
| 1 | 7 | 1.15E+05 | 8.50E+06 |
| 2 | 1 | 1.80E+05 | 8.43E+06 |
| 2 | 2 | 2.37E+05 | 7.33E+06 |
| 2 | 3 | 3.77E+05 | 6.53E+06 |
| 2 | 4 | 8.23E+04 | 5.40E+06 |
| 2 | 5 | 4.07E+05 | 7.53E+06 |
| 2 | 6 | 7.37E+05 | 1.93E+07 |
| 2 | 7 | 8.20E+05 | 5.77E+06 |

[A] cell titer is given in CFU per 1 g of rhizosphere sample

**Table S3 Statistics after demultiplexing and filtering.**

| Sample name | Number of reads | Yield in Mbps | Average quality |
|---|---|---|---|
| P482_maize_1 | 30925123 | 9066 | 35.7 |
| P482_maize_2 | 28740810 | 8431 | 35.66 |
| P482_maize_3 | 29182676 | 8600 | 35.62 |
| P482_tomato_1 | 24442543 | 7283 | 35.8 |
| P482_tomato_2 | 27623025 | 8162 | 35.63 |
| P482_tomato_3 | 31350329 | 9278 | 35.61 |
| P482_CTRL_1 | 25404161 | 7564 | 35.68 |
| P482_CTRL_2 | 28791672 | 8477 | 35.87 |
| P482_CTRL_3 | 20810455 | 5844 | 35.96 |

More information about the average quality score can be found at the Illumina website (https://www.illumina.com/science/education/sequencing-quality-scores.html)

**Table S4 RNAseq alignment statistics.**

| Sample name | Reference genome (Genbank)[A] | Filtered reads | Unique reads (%) | Multimapped reads (%) | Unmapped reads (%) | Reads per uniquely mapped (in millions) |
|---|---|---|---|---|---|---|
| P482_maize_1 | JHTS00000000.1 | 30925123 | 15.9 | 64.79 | 19.31 | 4.92 |
| P482_maize_2 | JHTS00000000.1 | 28740810 | 17.75 | 63.31 | 18.94 | 5.10 |
| P482_maize_3 | JHTS00000000.1 | 29182676 | 14.14 | 64.68 | 21.18 | 4.13 |
| P482_tomato_1 | JHTS00000000.1 | 24442543 | 5.3 | 72.93 | 21.77 | 1.30 |
| P482_tomato_2 | JHTS00000000.1 | 27623025 | 7.54 | 71.46 | 21 | 2.08 |
| P482_tomato_3 | JHTS00000000.1 | 31350329 | 9.36 | 68.9 | 21.74 | 2.93 |
| P482_CTRL_1 | JHTS00000000.1 | 25404161 | 15.74 | 64.23 | 20.03 | 4.00 |
| P482_CTRL_2 | JHTS00000000.1 | 28791672 | 15.77 | 65.73 | 18.5 | 4.54 |
| P482_CTRL_3 | JHTS00000000.1 | 20810455 | 19.13 | 67.56 | 13.31 | 3.98 |

[A] Reference number of sequences (contigs): 69. Reference number of bases: 5717769.

**Table S5 Tomato-specific genes od the differentiating response (GDRs) with the highest change in expression between.**

| Locus | Gene | $Log_2FC$ | Annotation |
|---|---|---|---|
| ***Upregulated*** | | | |
| BV82_3239 | *ytfE* | 6.36 | hemerythrin HHE cation binding domain protein |
| BV82_4743 | *hmp* | 6.13 | oxidoreductase NAD-binding domain protein |
| BV82_3240 | | 5.54 | putative membrane protein |
| BV82_3241 | *nnrS* | 4.99 | NnrS family protein |
| BV82_0044 | *cioA* | 4.92 | bacterial Cytochrome Ubiquinol Oxidase family protein |
| BV82_3243 | | 4.84 | putative dnrP protein |
| BV82_4439 | *metE* | 4.73 | 5-methyltetrahydropteroyltriglutamate-- homocysteine S-methyltransferase |
| BV82_0046 | | 4.50 | hypothetical protein |
| ***Downregulated*** | | | |
| BV82_3674 | | -4.48 | ABC transporter, substrate-binding, aliphatic sulfonates family protein |
| BV82_0910 | | -4.35 | hypothetical protein |
| BV82_1669 | *tauA* | -4.32 | taurine ABC transporter, a periplasmic binding protein |
| BV82_2326 | *trpB* | -4.28 | tryptophan synthase, beta subunit |
| BV82_2325 | *trpA* | -4.21 | tryptophan synthase, alpha subunit |
| BV82_2076 | | -3.97 | class II Aldolase and Adducin N-terminal domain protein |
| BV82_3637 | *ssuF* | -3.73 | molybdenum-pterin binding domain protein |
| BV82_3254 | | -3.67 | putative Lrp/AsnC family transcriptional regulator |

**Table S6 Maize-specific GDRs with the highest change in expression.**

| Locus | Gene | $Log_2FC$ | Annotation |
|---|---|---|---|
| ***Upregulated*** | | | |
| BV82_3988 | | 4.96 | hypothetical protein |
| BV82_2809 | | 4.20 | hypothetical protein |
| BV82_2032 | *mexE* | 4.12 | efflux transporter. RND family. MFP subunit |
| BV82_2904 | *copA* | 3.69 | copper-translocating P-type ATPase |
| BV82_4275 | *slyA1* | 3.25 | MarR family protein |
| BV82_2815 | *ripA_1* | 3.14 | bacterial regulatory helix-turn-helix s. AraC family protein |
| BV82_1378 | | 2.86 | bacterial regulatory s. TetR family protein |
| BV82_1235 | *nuoA* | 2.77 | NADH-ubiquinone/plastoquinone oxidoreductase. chain 3 family protein |
| ***Downregulated*** | | | |
| BV82_3753 | | -3.12 | TonB-dependent siderophore receptor |
| BV82_1217 | *furB* | -2.31 | $Fe^{2+}$ $Zn^{2+}$ uptake regulation protein |
| BV82_1856 | *polC1* | -2.04 | exonuclease family protein |
| BV82_1868 | *bioF* | -1.96 | 8-amino-7-oxononanoate synthase |
| BV82_0119 | *phrB* | -1.93 | FAD binding domain of DNA photolyase family protein |
| BV82_2744 | *thiC* | -1.90 | thiamine biosynthesis protein ThiC |
| BV82_0367 | *gcvH* | -1.63 | glycine cleavage system H protein |
| BV82_3597 | *rstB* | -1.51 | HAMP domain protein |

**Table S7 Genes of the shared response to exudates with the highest change in expression when compared to 1C medium without supplementation.**

| Locus | Gene | $Log_2FC$ Tomato | $Log_2FC$ Maize | Annotation |
|---|---|---|---|---|
| ***Upregulated*** | | | | |
| BV82_3873 | *arsC* | 2.42 | 2.66 | low molecular weight phosphotyrosine phosphatase family protein |
| BV82_3874 | *arsH* | 2.63 | 2.35 | arsenical resistance protein ArsH |
| BV82_2350 | *surf1* | 3.02 | 1.84 | SURF1 family protein |
| BV82_5016 | *fdnG* | 2.70 | 2.08 | formate dehydrogenase. alpha subunit |
| BV82_4419 | *bfr* | 1.87 | 2.67 | bacterioferritin |
| BV82_1741 | *tagT* | 2.21 | 2.16 | ABC transporter family protein |
| BV82_2984 | | 2.41 | 1.85 | plasmid replication region DNA-binding N-term family protein |
| BV82_1931 | *crp* | 2.03 | 2.05 | cyclic AMP receptor-like protein |
| ***Downregulated*** | | | | |
| BV82_0056 | | -6.83 | -6.84 | heme-binding A family protein |
| BV82_1668[A] | *tauB* | -4.52 | -1.87 | ABC transporter family protein |
| BV82_1666 | *tauD* | -4.11 | -3.83 | alpha-ketoglutarate-dependent taurine dioxygenase |
| BV82_1667 | *tauC* | -4.05 | -3.65 | binding--dependent transport system inner membrane component family protein |
| BV82_3001 | | -3.89 | -3.35 | hypothetical protein |
| BV82_0058 | *hasE* | -3.87 | -3.32 | type I secretion membrane fusion, HlyD family protein |
| BV82_0097 | | -3.66 | -4.31 | LysE type translocator family protein |
| BV82_3002 | *aroF* | -3.10 | -3.35 | 3-deoxy-7-phosphoheptulonate synthase (DAHP) |

[A] the difference in the mean expression of *tauB* in the presence of tomato and maize-derived compounds, although high (2.65 $log_2FC$), had $p=0.051$, therefore slightly below the assumed significance cutoff ($p<0.05$)

### Table S8 GC-MS and NMR quantification of tomato and maize exudates collected in water.

| Compound | Maize (ng/mg) | Tomato (ng/mg) | Ratio (Maize/ Tomato) |
|---|---|---|---|
| Glucose | 9947.00 | DNQ | - |
| trans-Aconitic acid | 134.56 | ND | - |
| Citric acid | 175.40 | 3.99 | 43.98 |
| Malic acid | 1481.02 | 88.13 | 16.80 |
| L-Leucine | 40.63 | 3.21 | 12.66 |
| L-Phenylalanine | 19.65 | 3.39 | 5.80 |
| Lactic acid | 92.63 | 48.62 | 1.91 |
| Glycolic acid | 8.80 | 4.94 | 1.78 |
| Myristic acid | 22.60 | 17.72 | 1.28 |
| Palmitic acid | 6.28 | 5.74 | 1.09 |
| Stearic acid | 8.28 | 8.70 | 0.95 |
| Succinic acid | 36.92 | 53.42 | 0.69 |
| Azelaic acid | 7.19 | 71.85 | 0.10 |
| Pimelic acid | 1.06 | 16.91 | 0.06 |
| Glycine | DNQ | DNQ | - |
| L-Tyrosine | DNQ | DNQ | - |
| Oleamide | DNQ | DNQ | - |

The quantity of compounds in given in ng per mg of exudate dry weight. Quantified by GC-MS using a mixture of standards, with the exceptions glucose, the latter quantified with NMR using lactic acid and malic acid as internal standards.
DNQ – detected not quantified (weak signal)
ND – not detected

### Table S9 Composition of standards for GC-MS

| **Compound** | **CAS no.** | **Supplier** | **Cat. no.** |
|---|---|---|---|
| ***Mix 10*** | | | |
| 5-Chloroindole-2-carboxylic acid | CAS 10517-21-2 | Alfa Aesar | A18626.03 |
| Azelaic acid | CAS 123-99-9 | Acros Organics | 401520250 |
| 4-Methoxyphenylacetic acid | CAS 104-01-8 | Acros Organics | 126021000 |
| Glycolic acid | CAS 79-14-1 | Acros Organics | 154510250 |
| **DL-Lactic acid** | CAS 50-21-5 | Acros Organics | 125060250 |
| Myristic acid | CAS 544-63-8 | Acros Organics | 156962500 |
| Palmitic acid | CAS 57-10-3 | Acros Organics | 129702500 |
| Pimelic acid | CAS 111-16-0 | Acros Organics | 131230250 |
| Stearic acid | CAS 57-11-4 | Alfa Aesar | A12244.06 |
| trans-Aconitic acid | CAS 4023-65-8 | Alfa Aesar | B20087.14 |
| ***Mix 6*** | | | |
| Citric acid | CAS 77-92-9 | Acros | A10395.30 |
| **DL-Lactic acid** | CAS 50-21-5 | Acros | 125060250 |
| L-Leucine | CAS 61-90-5 | Acros | 125121000 |
| L-Phenylalanine | CAS 63-91-2 | Acros | 130310250 |
| DL-Malic acid | CAS 6915-15-7 | Acros | 125252500 |
| Succinic acid | CAS 110-15-6 | Acros | 219552500 |

Both standards were prepared in concentration ranges from **0.2 to 0.0000002** mg $mL^{-1}$ and diluted 1:1 during silylation. Concentration of 0.002 was used to calculate the response of the detector for quantitative purposes. Mix10 was prepared and diluted in ethyl acetate, the solvent was evaporated in room temperature and the compounds were re-suspended in acetonitrile. To obtain Mix6, compounds were dissolved in acidic (~pH 2) ultrapure water, mixed, serially diluted, the solvent evaporated by freeze drying, with final re-suspension in acetonitrile to obtain the target concentrations.

**Table S10 Compound parameters of the MRM transitions in the LC-SRM method developed for the analysis of amino acids in maize and tomato root exudates.**

| Q1 Mass (Da) | Q3 Mass (Da) | MRM transition name | Transition type | DP (volts) | CE (volts) | CXP (volts) |
|---|---|---|---|---|---|---|
| 133.902 | 88.1 | Aspartic acid-1 | quantifier | 11 | 13 | 10 |
| 133.902 | 74 | Aspartic acid-2 | qualifier | 11 | 19 | 6 |
| 147.989 | 84 | Glutamic acid-1 | quantifier | 11 | 21 | 8 |
| 147.989 | 130.1 | Glutamic acid-2 | qualifier | 11 | 13 | 14 |
| 156.039 | 110 | Histidine-1 | quantifier | 56 | 19 | 18 |
| 156.039 | 83.1 | Histidine-2 | qualifier | 56 | 31 | 10 |
| 147.046 | 84.1 | Lysine-1 | quantifier | 21 | 19 | 8 |
| 147.046 | 130 | Lysine-2 | qualifier | 21 | 13 | 14 |
| 121.952 | 58.9 | Cysteine-1 | quantifier | 16 | 31 | 12 |
| 121.952 | 76 | Cysteine-2 | qualifier | 16 | 19 | 36 |
| 146.999 | 129.9 | Glutamine-1 | quantifier | 51 | 13 | 16 |
| 146.999 | 84 | Glutamine-2 | qualifier | 51 | 21 | 42 |
| 105.982 | 60 | Serine-1 | quantifier | 1 | 15 | 10 |
| 105.982 | 87.9 | Serine-2 | qualifier | 1 | 13 | 42 |
| 119.996 | 74 | Threonine-1 | quantifier | 11 | 15 | 34 |
| 119.996 | 102.1 | Threonine-2 | qualifier | 11 | 11 | 12 |
| 182.031 | 165.1 | Tyrosine-1 | quantifier | 21 | 13 | 10 |
| 182.031 | 136 | Tyrosine-2 | qualifier | 21 | 17 | 10 |
| 90.003 | 44.1 | Alanine-1 | quantifier | 1 | 13 | 20 |
| 90.003 | 45 | Alanine-2 | qualifier | 1 | 45 | 20 |
| 75.989 | 30 | Glicyne-1 | quantifier | 21 | 21 | 14 |
| 75.989 | 48 | Glicyne-2 | qualifier | 21 | 5 | 12 |
| 132.033 | 86 | Isoleucine-1 | quantifier | 31 | 13 | 16 |
| 132.033 | 69.2 | Isoleucine-2 | qualifier | 31 | 28 | 12 |
| 131.996 | 86 | Leucine-1 | quantifier | 51 | 13 | 16 |
| 131.996 | 44.1 | Leucine-2 | qualifier | 51 | 27 | 20 |
| 150.02 | 133 | Methionine-1 | quantifier | 56 | 13 | 6 |
| 150.02 | 104.2 | Methionine-2 | qualifier | 56 | 13 | 18 |
| 165.966 | 119.9 | Phenylalanine-1 | quantifier | 41 | 17 | 16 |
| 165.966 | 102.9 | Phenylalanine-2 | qualifier | 41 | 35 | 10 |
| 116.022 | 70.2 | Proline-1 | quantifier | 1 | 19 | 8 |
| 116.022 | 43.1 | Proline-2 | qualifier | 1 | 37 | 4 |
| 205.045 | 188 | Tryptophan-1 | quantifier | 1 | 13 | 10 |
| 205.045 | 146.1 | Tryptophan-2 | qualifier | 1 | 23 | 6 |
| 118.037 | 71.9 | Valine-1 | quantifier | 41 | 15 | 10 |
| 118.037 | 55.2 | Valine-2 | qualifier | 41 | 25 | 8 |
| 126 | 108 | Taurine-1 | quantifier | 25 | 15 | 15 |
| 126 | 43.9 | Taurine-2 | qualifier | 25 | 28 | 7 |
| 133.1 | 70 | Ornithine-1 | quantifier | 10 | 25 | 7 |
| 133.1 | 116 | Ornithine-2 | qualifier | 10 | 12.7 | 6 |
| 135.1 | 88.9 | Leucine-D3-1 | quantifier | 15 | 13 | 11 |
| 135.1 | 45.3 | Leucine-D3-2 | qualifier | 15 | 27 | 9 |
| 175.034 | 70.1 | Arginine-1 | quantifier | 41 | 29 | 8 |
| 175.034 | 116.1 | Arginine-2 | qualifier | 41 | 19 | 8 |

**Table S11 The calculated $log_2$ fold differences in the quantity of 21 amino acids in maize root exudates comparing to tomato root exudates, with their corresponding p-values of statistical significance.**

| Amino acid | log2 fold difference[A] | p-value | Statistical significance ($p<0.05$) |
|---|---|---|---|
| Ala | 1,6542 | 1,14 E-02 | yes |
| Arg | 2,5996 | 2,05 E-06 | yes |
| Asp | 0,9521 | 6,49 E-02 | no |
| Cys | -1,5467 | 1,90 E-01 | no |
| Gln | -0,2455 | 7,51 E-01 | no |
| Glu | -1,7185 | 8,94 E-02 | no |
| Gly | -0,2519 | 8,29 E-01 | no |
| His | 2,3553 | 5,66 E-02 | no |
| Ile | 0,9354 | 6,30 E-02 | no |
| Leu | 2,2029 | 1,58 E-02 | yes |
| Lys | 0,9745 | 2,14 E-03 | yes |
| Met | 0,5612 | 1,53 E-01 | no |
| Orn | 1,4055 | 5,23 E-07 | yes |
| Phe | 2,1967 | 5,54 E-03 | yes |
| Pro | 2,7591 | 9,17 E-11 | yes |
| Ser | -0,5485 | 5,61 E-01 | no |
| Tau | -3,1630 | 3,00 E-10 | yes |
| Thr | 1,9070 | 2,28 E-02 | yes |
| Trp | 1,2491 | 8,15 E-04 | yes |
| Tyr | 2,7140 | 5,05 E-08 | yes |
| Val | 2,3742 | 2,23 E-05 | yes |

[A] tomato exudate were used as the reference sample.
